# Chemical-Genetic Interrogation of Nuclear Size Control Reveals Cancer-Specific Effects on Cell Migration and Invasion

**DOI:** 10.1101/2020.01.10.902148

**Authors:** Andrea Rizzotto, Sylvain Tollis, Nhan T. Pham, Jan Wildenhain, Nikolaj Zuleger, Jeremy T. Keys, Dzmitry Batrakou, Jayne Culley, Sarah Zheng, Jan Lammerding, Neil O. Carragher, Valerie G. Brunton, Manfred Auer, Mike Tyers, Eric C. Schirmer

**Affiliations:** The Institute of Cell Biology, University of Edinburgh, Edinburgh, EH9 3BF, UK; Institute for Research in Immunology and Cancer, Université de Montréal, Montréal, Québec, H3T 1J4, Canada; Institute of Quantitative Biology, Biochemistry and Biotechnology, University of Edinburgh, Edinburgh, EH9 3BF, UK; Nancy E. and Peter C. Meinig School of Biomedical Engineering & Weill Institute for Cell and Molecular Biology, Cornell University, Ithaca, New York, 14853, USA; Edinburgh Cancer Research UK Centre, Institute of Genetics and Molecular Medicine, University of Edinburgh, Edinburgh EH4 2XR

## Abstract

Lower survival rates for many cancer types correlate with increases or decreases in nuclear size/scaling in a tumor-type/tissue-specific manner. Postulating that nuclear size changes confer a fitness advantage on tumor cells, we screened for FDA/EMA-approved compounds that reverse tumor nuclear size changes in cell lines from three such tumor types: prostate adenocarcinoma, colonic adenocarcinoma, and small-cell squamous lung cancer. We found distinct, largely non-overlapping sets of compounds that either rectify or exacerbate nuclear size changes for each tumor type. Nuclear size phenotypes across cell lines clustered particular classes of compounds including serotonin uptake inhibitors, cyclo-oxygenase inhibitors, beta-adrenergic receptor agonists, monoamine oxidase inhibitors, and Na^+^/K^+^ ATPase inhibitors. Nearly all compounds selected for further investigation inhibited cell migration and/or invasion, suggesting that targeting nuclear size control pathways in chemotherapy regimens could improve patient survival.

## Introduction

Nuclear size changes have been used in cancer diagnosis since the 1860s^1^ and are still used for diagnosis and prognostic grading of later stage, higher-grade tumors^2^. Characteristic nuclear size changes observed are independent of ploidy for at least a dozen cancer types, suggesting a primary dysfunction in regulation of size/scaling in the cell. However, it remains unclear whether nuclear size alterations contribute to increased metastasis, in part because the directionality and degree of size changes is tumor/tissue-type specific^3, 4^. For example, in small-cell squamous lung cancer and osteosarcoma smaller nuclear size correlates with increased metastasis^5, 6^ while for breast, prostate, colon and several other cancer types increased nuclear size correlates with increased metastasis^7,8,9,10^.

While absolute nuclear size changes are used in current diagnostics, in some tumor types the size change is associated with disruption of cell scaling^11^. In normal cells and tissues, the nuclear-to-cytoplasm (N/C) ratio, also called the karyoplasmic ratio, is maintained during the cell cycle^12, 13^, throughout which the nucleus increases several-fold in volume^14^. In contrast, the N/C ratio is altered in cancer cells by non-correlated changes in nuclear size and/or cell size. It has long been known that a general scaling mechanism for maintaining the N/C ratio is conserved from higher eukaryotes to yeast^15, 16^ but the mechanistic basis for this scaling remains uncertain. A number of proteins have been implicated in regulating absolute nuclear size and N/C scaling. Nuclear size is influenced by nuclear envelope (NE) proteins, such as the lamins that form the nucleoskeleton^17^ and the outer nuclear membrane nesprins that connect to the cytoskeleton^18^. The inner nuclear membrane protein LEM2 also regulates nuclear size by controlling membrane flow^19^, possibly through interactions with lamins, chromatin^20, 21^ and protein kinases ^22^. Recent genetic screens in yeast have implicated nucleocytoplasmic transport, LINC complexes, and RNA processing/splicing in nuclear size and N/C ratio control^23, 24, 25^.

As nuclear size and/or N/C ratio scaling is disrupted in cancer cells with increased metastatic potential, we reasoned that these changes might contribute an advantage to the tumor by facilitating metastasis. From a regulatory and structural perspective, the NE connects to both the genome and the cytoskeleton. Dysregulation of nuclear size could therefore yield metastatic advantages ranging from altered gene regulation, nuclear signalling and/or nucleo-cytoplasmic transport to mechanical effects enabling faster migration and/or easier invasion of cancer cells through epithelial barriers. Indeed, disruption of laminA reduces cell migration^26^. Moreover, mechanical forces that would be expected to be altered by nuclear size changes are important for the progress of metastasis^27^ and cellular traction stresses are linked to metastatic potential^28^. For these reasons, we speculated that restoring nuclear size to a more normal range might reduce metastatic potential.

As each tumor/tissue type has different characteristic size/scaling changes, and because many parameters likely affect nuclear size, it is difficult to correlate cancer genome sequence information with alterations in nuclear size control. Here, we used a chemical-genetic approach to systematically probe the pathways that affect nuclear size in cancer cells. We screened for FDA/EMA-approved compounds that rectify nuclear size in three cancer cell lines derived from different tissues: two in which nuclear size increases correlate with worse grade (prostate, colon adenocarcinoma) and one in which nuclear size decrease correlates with worse grade (lung). The majority of compounds identified had tumor/tissue/cell line-specific effects on nuclear size or scaling, typically only affecting one of the three cancer cell lines. Clustering compounds according to their therapeutic classes/mechanism of action identified many compound classes with characteristic nuclear size phenotypic signatures across cell lines and conditions. Among these, some altered the absolute nuclear size but not N/C ratio or vice-versa, suggesting different modes of action. Detailed investigation of five compounds revealed cell line-specific effects on cell invasion, consistent with nuclear size changes observed. These results suggest that pharmacological restoration of nuclear size control represents a new strategy to combat metastatic cancer.

## Results

### Nuclear size/scaling chemical screen

We screened for compounds that alter nuclear size and/or N/C ratio in three tumor types using a microscopy-based approach (Fig. 1). PC3 and HCT116 cells respectively represented late-stage prostate cancer and colonic adenocarcinoma where nuclear size increases compared to the healthy tissues reflect worse cancer grades. H1299 cells represented small-cell squamous lung carcinoma where decreased nuclear size correlates with a worse grade. Cells plated onto 96-well optical plates were treated for 6 h or 36 h with the Prestwick chemical library of 1,120 previously approved drugs (PAD, 2015 version), all at a concentration of 10 µM. We chose the 6 h time-point to reveal compounds that did not require post-mitotic NE re-assembly to elicit size changes and the 36 h timepoint to identify compounds with low toxicity. To align with standard cancer diagnostic procedures, we used nuclear/cell area from imaging (focal) cross sections as our core size metric, as opposed to 3D reconstruction of nuclear/cell volume. Nuclear area was monitored with stably expressed H2B-mRFP and total cell area from CellMask DeepRed cytosol staining (Fig. 1a). 350-1,000 cells/condition were imaged using a Perkin-Elmer OPERA^TM^ confocal microscope for each screen. Two full replicate screens were undertaken for each cell line and treatment duration, based on reproducibility in a pilot screen using three replicates (Methods; Supplementary Figure S1; Supplementary Table S1).

**Fig. 1.**
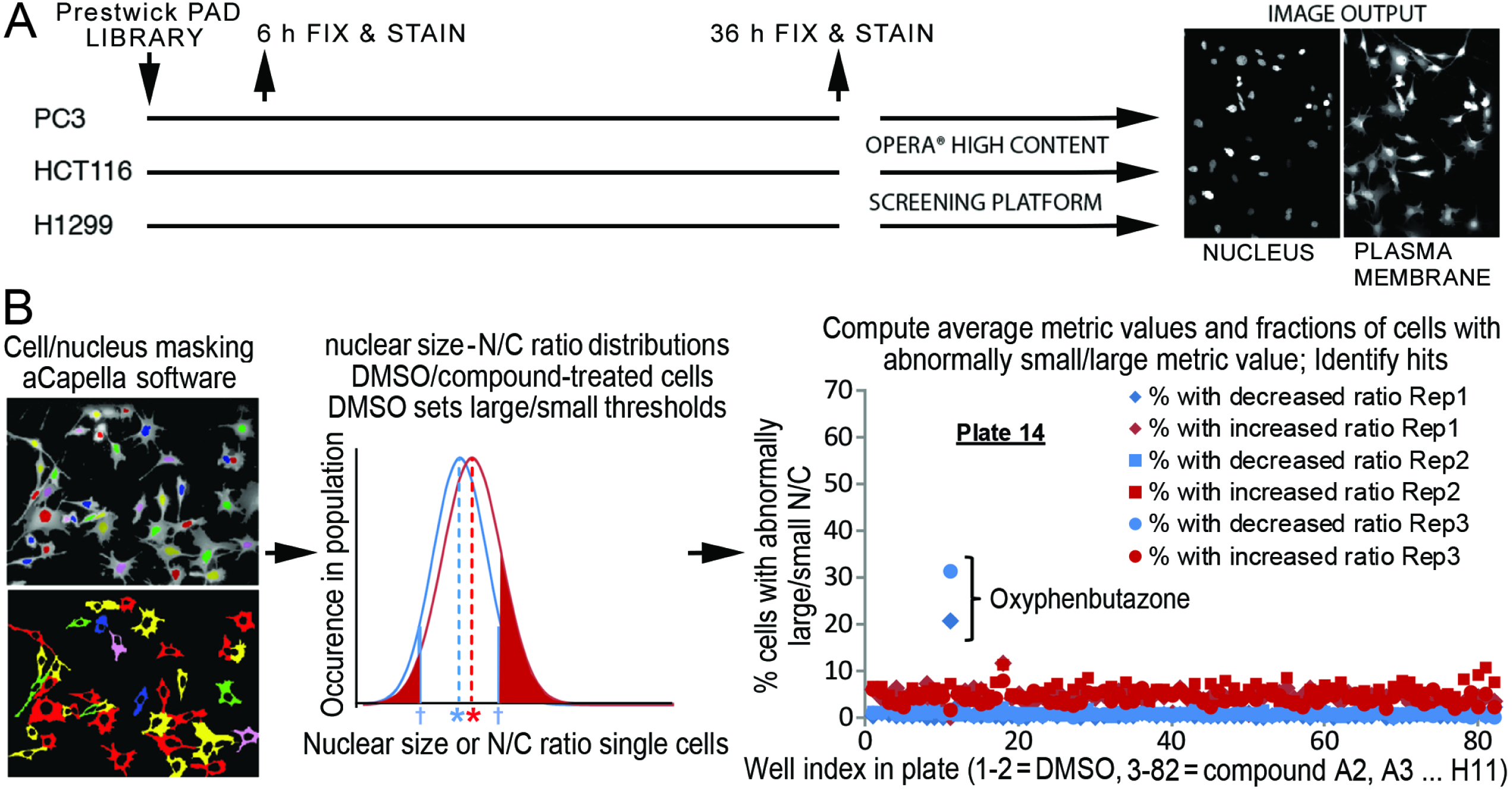
High-throughput screen for compounds affecting nuclear size. **a** Cell growth, treatment and imaging methodology. Compounds were added to cells pre-grown for 24 h on imaging plates, and cells were fixed and stained after 6 or 36 h of treatment. 350-1,000 cells per screen per condition were imaged on the OPERA high-content screening instrument. **b** Data analysis methodologies. Within each well, individual cells (left, bottom image) and their nuclei (left, top image) were masked respectively based on CellMaskDeepRed cytosol staining and H2B-mRFP staining using the Acapella^®^ built-in analysis software. For nuclear and N/C ratio metrics, well-based distributions of individual cell metric values were computed for DMSO (blue) and compound-treated wells (red, middle plot). The former was used to define outlier thresholds (blue crosses, see Methods), from which the fractions of outlier cells (red-shaded regions) were computed for compound-treated wells (shown regions) and DMSO-treated cells (not shown). Metrics averaged within each well were also computed (see blue and red asterisks for DMSO- and compound-treated wells, respectively). For each cell line/time point, data was collected from all wells across the library/replicates (right plot, example shown is plate 14 for PC3, 6 h). Standard statistical analyses of each dataset was used to identify compounds (e.g., oxyphenbutazone) that increased the fractions of outlier cells with either small/large nucleus and/or N/C ratio beyond non-specific effects of DMSO and inactive compounds. Compounds for which fractions of outlier cells exceeded the distribution across the library were defined as hits.

Two complementary quantification strategies, one average-based and the other outlier-based, were used to analyse effects of compounds on both the absolute nuclear size and the N/C ratio (i.e., relative nuclear size). Hence, we performed four separate analyses of the dataset. Average-based analyses are traditionally used by cytologists to grade tumors^29, 30^. To implement this approach, the average values of the size metrics were determined across all imaged cells for each compound well and in-plate DMSO-treated controls. However, shifting the average metric values up or down does not obligatorily reflect loss of size regulation, but could reflect a change in the cell-regulated value to which the metric is set. Thus, to detect compounds that specifically disrupt nuclear size control by increasing the metric variance across a cell population, we also analysed the same data using an outlier-based approach. We computed for each well the fractions of cells with abnormally large or small metric value (i.e., outlier cells). The outlier metric value refers to values beyond the outlier thresholds defined using standard procedures with respect to the distribution of metric values for all in-plate DMSO control wells (see Fig. 1B and Methods). For the average-based and outlier-based methods, respectively, the average metric values and the fractions of outlier cells from all wells were analysed statistically to identify hit compounds that perturb the metric beyond the typical metric variability across the library. This approach should reduce non-specific compound effects on nuclear size (see Methods). Hit compounds were confirmed using direct comparison with in-plate DMSO controls in Wilcoxon rank statistical tests.

Strikingly, we observed little overlap in the sets of compounds affecting absolute nuclear size between the different cell lines using average-based analysis (Fig. 2a; Table 1; Supplementary Table S2). The combined short (6 h) and long (36 h) time-points yielded a total of 112 compound hits that altered mean nuclear size in PC3 cells (32 increase, 80 decrease), 105 hits in HCT116 cells (35 increase, 70 decrease) and 111 hits in H1299 cells (70 increase, 41 decrease). Compounds with the strongest functional effects are listed in Table 1. Different compounds reversed cancer-associated nuclear size changes either in a cell line-specific or non-specific manner (Table 1, Supplementary Figure S2). Cell types in which worse grade correlated with increased nuclear size (PC3, HCT116) had more active compounds at the long treatment time while the cell type in which smaller nuclear size correlated with worse grade (H1299) yielded more active compounds at the short treatment time (Fig. 2b; Table 1; Supplementary Table S2). It is possible that long term effects on nuclear size may indicate a requirement for restructuring the NE in mitosis, while selective short-term effects may indicate homeostatic adaptation to perturbation.

**Fig. 2.**
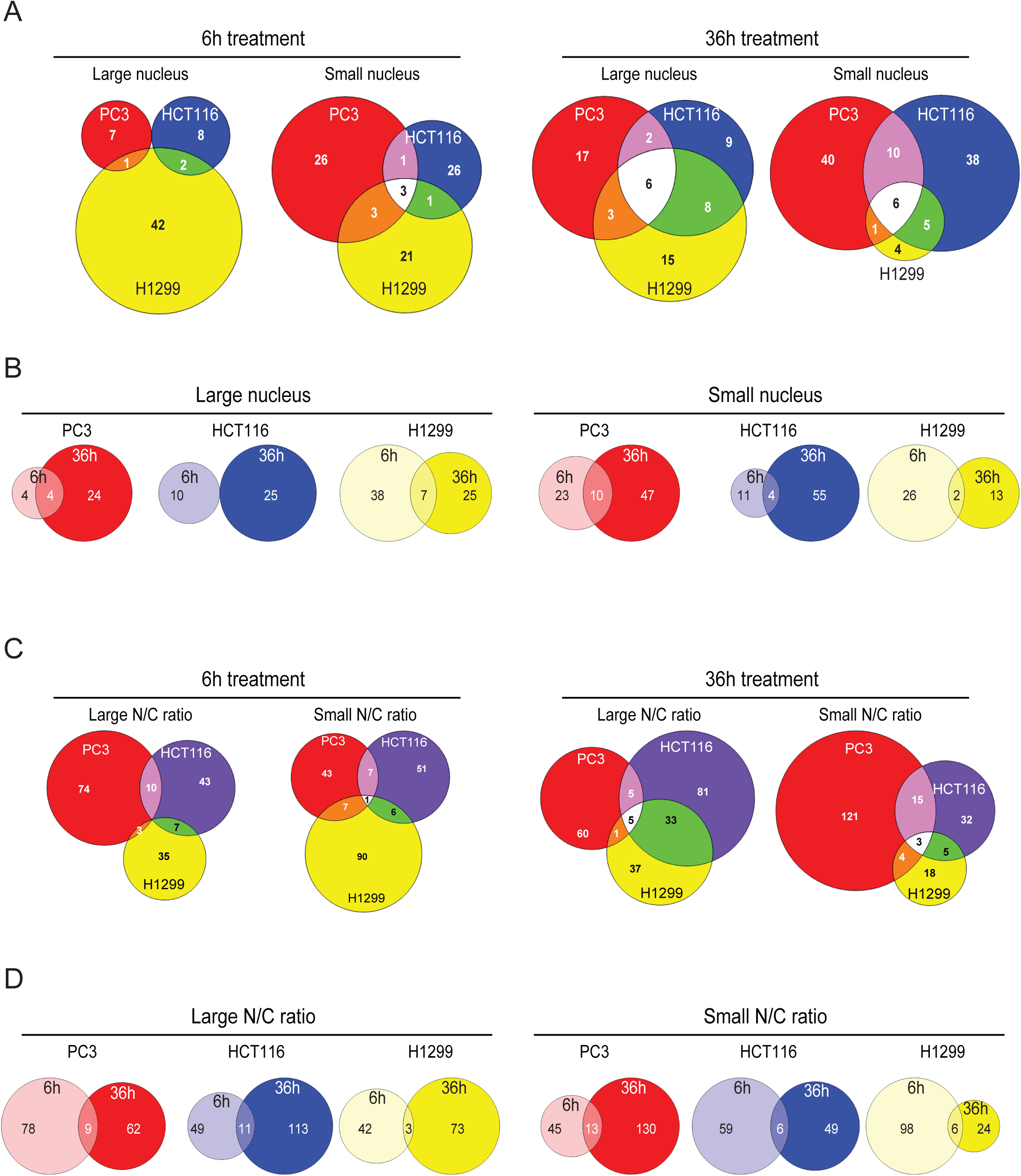
Tumor/tissue-type specific effect of NSR compounds. **a** Venn diagrams showing the overlap in compounds increasing or reducing mean absolute nuclear size in PC3 (red), HCT116 (blue) and H1299 (yellow) tumor cell types treated for 6h (left) and 36 h (right). **b** Venn diagrams showing the overlap in compounds increasing (left three diagrams) or reducing (right three diagrams) mean nuclear size upon 6h (light colours) and 36 h (dark colours) treatment. Shown are PC3 (red), HCT116 (blue) and H1299 (yellow) tumor cell types. **c** Venn diagrams showing the overlap in compounds increasing or reducing mean N/C ratio in PC3 (red), HCT116 (blue) and H1299 (yellow) tumor cell types treated for 6h (left) and 36 h (right). **d** Venn diagrams showing the overlap in compounds increasing (left three diagrams) or reducing (right three diagrams) mean nuclear size upon 6h (light colours) and 36 h (dark colours) treatment. Shown are PC3 (red), HCT116 (blue) and H1299 (yellow) tumor cell types.

**Table 1.** NSR compounds that correct cancer-related nuclear size or N/C ratio changes in one or several of the three tumor types beyond detection thresholds in at least 2 replicates. Compound classes currently used in chemotherapy (e.g., microtubule inhibitors, DNA intercaling agents) have been removed from this table and are discussed in Supplementary Figure S3. Compound classes with consistent phenotypes across the class (e.g., beta adrenergic receptor agonists, Na+/K+ ATPase inhibitors) are described in Figure 5.

| **Compound – therapeutic group** | **Nuclear change correction** | **Strength, Replicates** | **Nuclear change aggravation** |
| --- | --- | --- | --- |
| Disulfiram – Dopamine beta hydrolase inhibitor | PC3, N, 6 h<br>PC3, N/C, 36 h<br>H1299, N/C, 36 h | Mild, 2/3<br>Strong, 2/2<br>Strong, 2/2 |  |
| Tomatine – cholinergic inhibition | PC3, N&N/C, 6 h<br>PC3, N&N/C, 36 h<br>HCT116, N, 6 h<br>HCT116, N, 36 h | Strong, 3/3<br>Strong, 1/2<br>Strong, 1/2<br>Strong, 1/2 |  |
| Astemizole – Histamine antagonist | PC3, N&N/C, 6 h<br>PC3, N&N/C, 36 h | Mild, 3/3<br>Strong, 2/2 | H1299, N/C, 6-36 h |
| Perhexiline maleate – Ca<sup>2+</sup> channel blocker | PC3, N, 6 h<br>PC3, N, 36 h<br>HCT116, N, 36 h | Strong, 2/3<br>Strong, 2/2<br>Mild, 1/2 |  |
| Ebselen – Cox inhibitor | PC3, N, 6 h<br>PC3, N/C, 6 h | Mild, 3/3<br>Strong, 3/3 |  |
| Alexidine dihydrochloride - Detergent | PC3, N&N/C, 6 h<br>PC3, N&N/C, 36 h<br>HCT116, N&N/C, 36 h<br>HCT116, N/C, 6 h<br>H1299, N/C, 6 h<br>H1299, N/C, 36 h, rep2 | Mild, 2/3<br>Strong, 2/2<br>Mild to strong, 2/2<br>Strong, 1/2<br>Strong 2/2 | H1299, N/C, 36 h, rep1 |
| Cantharidin – Serine protease releaser | PC3, N, 6 h<br>PC3, N, 36 h<br>HCT116, N, 36 h | Mild, 3/3<br>Mild, 2/2<br>Mild, 2/2 | HCT116, N, 6 h<br>H1299, N, 6 h<br>PC3, N/C, 6-36 h |
| Oxyphenbutazone – Cox inhibitor | PC3, N&N/C, 6 h<br>PC3, N&N/C, 36 h<br>HCT116, N, 6 h<br>HCT116, N, 36 h<br>H1299, N/C, 6 h<br>H1299, N/C, 36 h | Strong, 2/3<br>Strong, 2/2<br>Strong, 1/2<br>Strong, 2/2<br>Strong, 2/2<br>Strong, 1/2 | H1299, N, 6 h<br>H1299, N, 36 h |
| Anisomycin – Acetylcholine esterase inhibitor | PC3, N, 36 h<br>H1299, N&N/C, 36 h | Strong, 2/2<br>Strong, 2/2 | Possibly<br>HCT116, N/C |
| Roxatidine acetate – Histamine antagonist | PC3, N, 36 h<br>H1299, N, 6 h<br>H1299, N/C, 36 h | Strong, 2/2<br>Mild, 2/2<br>Strong, 2/2 | PC3, N, 6 h |
| Metixene HCl – Cholinergic inhibition | PC3, N, 36 h | Mild, 2/2 | H1299, N, 6 h |
| Parthenolide –MAO inhibitor | PC3, N, 36 h<br>H1299, N, 36 h | Mild, 2/2<br>Mild, 2/2 | HCT116, N/C, 36 h |
| Prenylamine lactate – Ca<sup>2+</sup> channel stimulation | PC3, N&N/C, 36 h | Strong, 2/2 |  |
| Emetine dihydrochloride – Protein synthesis inhibitor | PC3, N, 36 h<br>H1299, N, 6 h | Strong, 2/2<br>Mild, 2/2 | H1299, N, 36 h |
| Piperlongumine - alkaloid | PC3, N, 36 h<br>H1299, N/C, 36 h | Strong, 2/2<br>Strong, 2/2 | PC3, N/C, 6 h<br>HCT116, N/C, 6 h |
| Lycorine HCl – protein translation inhibitor | PC3, N, 36 h<br>HCT116, N, 36 h | Mild, 2/2<br>Mild, 2/2 | PC3, N/C, 6 h |
| Sertraline – 5HT uptake inhibitor | PC3, N, 6 h<br>PC3, N, 36 h<br>HCT116, N, 36 h<br>H1299, N/C, 6 h | Mild, 3/3<br>Strong, 2/2<br>Strong, 1/2<br>Mild, 2/2 | PC3, N/C, 6 h<br>H1299, N, 6 h |
| Danazol – Hormone estrogen antagonist | HCT116, N, 36 h | Mild, 2/2 | PC3, N/C, 6 h |
| Griseofulvin – antifungal drug | HCT116, N, 36 h | Mild to Strong, 2/2 |  |
| Puromycin dihydrochloride - antimetabolite | HCT116, N, 36 h<br>H1299, N, 6 h | Strong, 2/2<br>Strong, 2/2 | H1299, N, 36 h |
| Resveratrol – Cox inhibitor | PC3, N, 36 h<br>HCT116, N, 36 h | Mild, 2/2<br>Strong, 2/2 | H1299, N, 36 h<br>PC3, N/C, 6h |
| Cilostazol – phosphodiesterase inhibitor | HCT116, N, 36 h | Strong, 2/2 | PC3, N/C, 6 h |
| Eburnamonine - Alkaloid | HCT116, N, 36 h | Mild to Strong – 2/2 |  |
| Monensin sodium salt – membrane ionophores producer | HCT116, N, 36 h | Strong, 2/2 | H1299, N, 36 h |
| Dienestrol – Non-steroidal estrogen | HCT116, N, 36 h | Mild – 2/2 |  |
| Azacytidine-5-antimetabolite | HCT116, N, 36 h | Mild – 2/2 | H1299, N, 36 h |
| Hydroflumethiazide – Na<sup>+</sup> Cl<sup>-</sup> transport inhibitor | H1299, N, 6 h | Strong 2/2 |  |
| Metaraminol bitartrate – Alpha adrenergic agonist | H1299, N&N/C, 6 h | Strong 2/2 |  |
| Flufenamic acid – Cox inhibitor | H1299, N, 6 h | Strong 2/2 |  |
| Trimethoprim – Folic acid antagonist | H1299, N, 6 h | Mild, 2/2 |  |
| Fenspiride HCl – Bradykinin antagonist | H1299, N, 6 h | Strong 2/2 |  |
| Methylprednisolone 6-alpha - Glucocorticoid | H1299, N, 6 h<br>HCT116, N/C, 36 h | Strong 2/2<br>Mild, 2/2 | HCT116, N, 36 h |
| Debrisoquin sulfate – Catecholamine depletor | H1299, N, 6 h<br>H1299, N, 36 h | Strong, 2/2<br>Mild, 2/2 |  |
| Harmol HCl monohydrate - anxiolytic | H1299, N, 6 h | Strong, 2/2 |  |
| Methoxy-6-harmalan – Benzodiazepine receptor ligand | H1299, N, 6 h<br>HCT116, N/C, 36 h | Mild, 2/2<br>Strong, 1/2 |  |
| Levonordefrin – alpha adrenergic agonist | H1299, N, 6 h<br>HCT116, N/C, 36 h | Strong, 2/2<br>Strong, 2/2 | HCT116, N, 36 h |
| N-Acetyl-L-leucine – anti-vertigo drug | H1299, N, 6 h | Mild, 2/2 |  |
| (+)-Isoproterenol (+)-bitartrate salt - Bronchodilator | H1299, N, 6 h<br>HCT116, N/C, 6 h | Strong ,2/2<br>Strong, 2/2 |  |
| Proscillaridin A – cardiac glycoside | H1299, N, 6 h<br>H1299, N, 36 h | Strong 2/2<br>Strong 1/2 |  |
| Etilefrine HCl – alpha adrenergic agonist | H1299, N, 6 h<br>HCT116, N/C, 6 h | Strong 2/2<br>Strong, 2/2 |  |
| Letrozole – P450 inhibitor | H1299, N, 6 h<br>HCT116, N/C, 6 h | Strong 2/2<br>Strong, 2/2 |  |
| (+,-)-Synephrine – alpha adrenergic agonist | H1299, N, 6 h<br>HCT116, N/C, 6 h | Mild 2/2<br>Mild, 2/2 |  |
| Verteporfin – treatment of ARMD | H1299, N, 6 h | Mild, 2/2 |  |
| Monobenzone – melanin inhibition | H1299, N, 36 h | Strong 2/2 | HCT116, N, 36 h |
| Trifluridine – DNA replication inhibitor | H1299, N, 36 h<br>PC3, N/C, 36 h | Strong 2/2<br>Strong, 2/2 | PC3, N, 6 h<br>HCT116, N, 36 h |
| Antimycin A – mitochondrial ETC inhibitor | PC3, N/C, 6 h | Mild 2/3 |  |
| Mitoxantrone dihydrochloride – topoisomerase II inhibitor | H1299, N/C<br>PC3, N/C, 36 h | Strong ½<br>Mild, 2/2 | HCT116, N/C, 36 h |
| Paroxetine HCl – 5HT uptake inhibitor | PC3, N, 6 h | Mild, 2/3 | PC3, N/C, 6 h |
| Parbendazole - Anthelmintic | HCT116, small N outliers, 36 h<br>H1299, large& small Noutliers, 36 h | Mild, 2/2<br>Mild, 2/2 | PC3, large N outliers, 6 h |
| Camptothecine (S,+) – Topoisomerase I inhibitor | All lines, large&small N outliers, 36 h | Mild to strong, 2/2 |  |

There was a strong overlap between compound sets identified using average-based versus outlier-based analyses for absolute nuclear size although the outlier-based method tended to identify more hits (Supplementary Figure S2a, Table 1, compare Supplementary Tables S2, S4). The convergence of two fundamentally different analysis methods underscores the robustness of our data. At the same time, differences between the two analyses might be biologically relevant. In particular, average-based only hits might alter the nuclear size set point but not homeostatic control whereas outlier-based only hits might perturb homeostasis but not the nuclear size set point.

We carried out analogous average- and outlier-based analyses of the N/C ratio, which normalizes nuclear size changes relative to overall cell size. Both analyses identified cancer-type specific compound sets, with many compounds the same as those identified by the absolute nuclear size analyses (Fig. 2c-d; Table 1; Supplementary Tables S3, S5). However, N/C ratio analyses identified many additional compounds. Combining short and long time-points for the average analysis yielded a total of 237 hits for PC cells (149 increase, 188 decrease), 287 hits in HCT116 cells (173 increase, 114 decrease) and 246 hits in H1299 cells (118 increase, 128 decrease). That more than twice as many compounds were identified by the N/C ratio analysis than absolute nuclear size indicates that many compounds can also affect cell size.

In the following sections, unless otherwise specified, we used the absolute nuclear size metric and the average analysis methodology. This metric/methodology association gave the most reproducible results in our triplicate pilot screen (Supplementary Figure S1; Supplementary Table S1). We used comparison of absolute and relative nuclear size metrics across time-points to provide mechanistic information on whether the compounds act in a cell size-dependent or -independent fashion. We used comparison of average and outlier analyses to discover compounds that alter nuclear size setpoint *per se* as opposed to

### General versus cell line-specific compound classes

All compounds were sorted according to pharmacological class/mechanism-of-action and clustered according to increased (Fig. 3) and decreased (Fig. 4) nuclear size phenotypes. Strikingly, certain compound classes showed phenotypic clustering across cell lines. DNA intercalating agents increased absolute nuclear size (but less so the N/C ratio) in all three cell lines (Supplementary Figure S3a), with strongest effect at the 36 h timepoint. Microtubule polymerisation inhibitors similarly tended to increase the fraction of cells with abnormally large nuclei across all cell lines, also with enrichment at the longer timepoint (Supplementary Figure S3b). However, microtubule poisons also increased the fraction of cells with abnormally small nuclei, such that the overall effects on average nuclear size were subtle. In particular, microtubule poisons increased the fraction of PC3 cells with abnormally large N/C ratios at 6 h without affecting the proportion of cells with large nuclear size (Supplementary Figure S3c, compare bottom and top heat maps). In contrast, by 36 h the initial increase in N/C ratio disappeared while nuclear size was now strongly affected in both directions. Microtubule poisons such as fenbendazole, colchicine, nocodazole, paclitaxel, albendazole, and podophyllotoxin may initially increase nuclear size, but the cell subsequently adjusts its cytoplasm to compensate. These results illustrate that multiparametric analyses can reveal size and scaling defects that would be missed by monitoring only a single parameter. As both DNA intercalating agents and microtubule polymerisation inhibitors are mainstays of chemotherapy regimens, these findings also exemplify the utility of the nuclear size screens in identifying general anti-cancer compounds. Indeed, our screens recovered most known antineoplastic compounds present in the Prestwick library, with variable effects on nuclear size (Supplementary Figure S3d). However, as such chemotherapeutic agents are generally toxic we focused on compounds with cell-type specific effects that may circumvent this drawback.

**Fig. 3.**
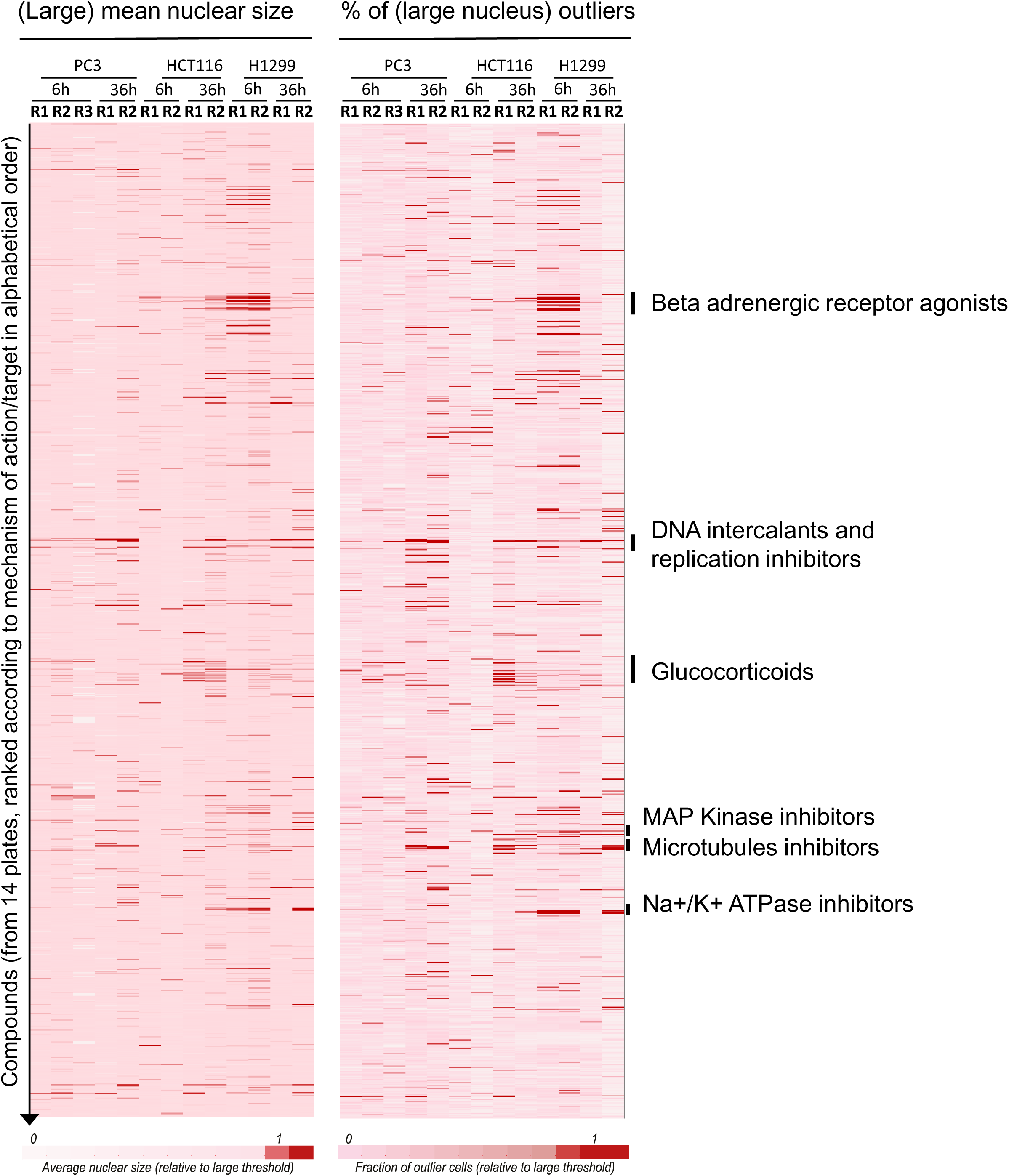
NSR compounds that increase nuclear size. Heat maps showing the mean nuclear size (left) and the mean N/C ratio (right) across the compounds collection (vertical) and across tumor type/replicates (horizontal). Nuclear size and N/C ratio were normalized to the detection thresholds for large nuclei (see Methods), and colour-coded as indicated, where a darker red corresponds to a stronger phenotype (larger nucleus or N/C ratio). Compounds were ranked by mechanism of action/therapeutic class, in alphabetic order. Particular classes of compounds showing interesting phenotypes are indicated.

**Fig. 4.**
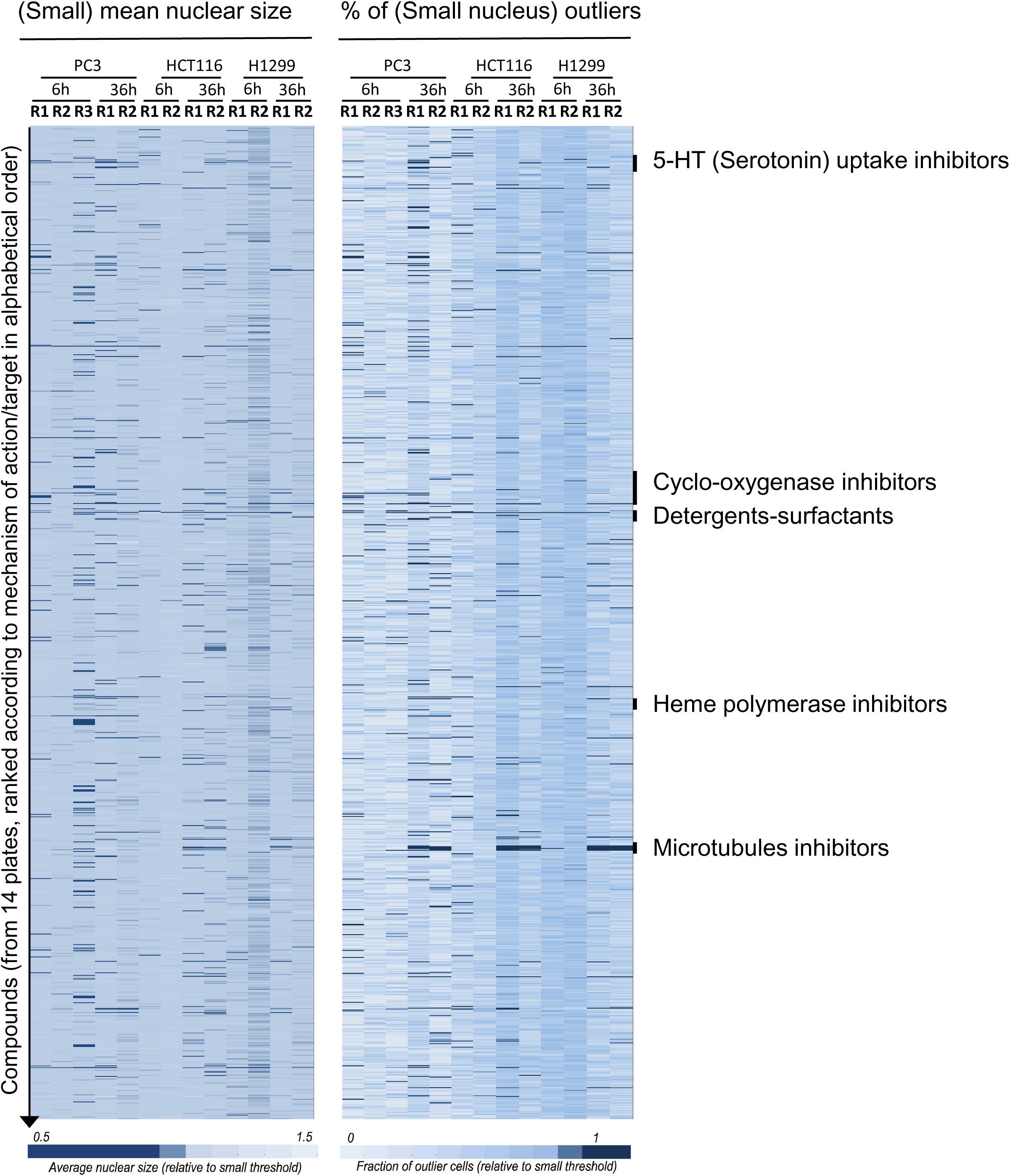
NSR Compounds that decrease nuclear size. Heat maps showing the mean nuclear size (left) and the mean N/C ratio (right) across the compounds collection (vertical) and across tumor type/replicates (horizontal). Nuclear size and N/C ratio were normalized to the detection thresholds for small nucleus (see Methods), and colour-coded as indicated, where a darker blue corresponds to a stronger phenotype (smaller nucleus or N/C ratio). Note that the data shown is the same as data shown on Figure 3, but normalised to small size thresholds and colour-coded in reverse (darker blue: smaller nucleus or N/C ratio). Compounds were ranked by mechanism of action/therapeutic class, in alphabetic order. Particular classes of compounds showing interesting phenotypes are indicated.

As nuclear size changes can occur in both directions, we refer to any compound that reversed a cancer-associated nuclear size phenotype as a nuclear size rectifier (NSR), whether it acts to make a large nucleus smaller or a small nucleus larger. Several compound classes were NSRs in only one or two of the three cancer cell lines (Fig. 5). Beta-adrenergic receptor agonists (BAAs, Fig. 5a) such as salbutamol, fenoterol, clenbuterol, isoetharine mesylate, and isoproterenol, but not antagonists such as propafenone, levobunolol, and betaxolol (Supplementary Figure S4a), reduced the nuclear size of HCT116 cells and increased the nuclear size of H1299 cells, i.e., towards a more normal size in both cases. BAAs affected absolute nuclear size but not the N/C ratio in H1299 cells while affecting N/C ratios and not absolute nuclear size in HCT116 cells (compare Fig. 5a with Supplementary Figure S4a). This example illustrates that the same compound class can elicit distinct nuclear size changes in different cell lines/cancer types. In contrast, Na+/K+ ATPase inhibitors, such as digitoxigenin and lanatoside, selectively increased nuclear size only in H1299 cells (Fig. 5a) and thus may exhibit selectivity towards lung cancer. Similarly, glucocorticoids increased absolute nuclear size and to a lesser extent the N/C ratio specifically in HCT116 cells (Supplementary Figure S4b). The widespread use of glucocorticoids in colon cancer therapy^31^ may thus engender unintended adverse characteristics.

**Fig. 5.**
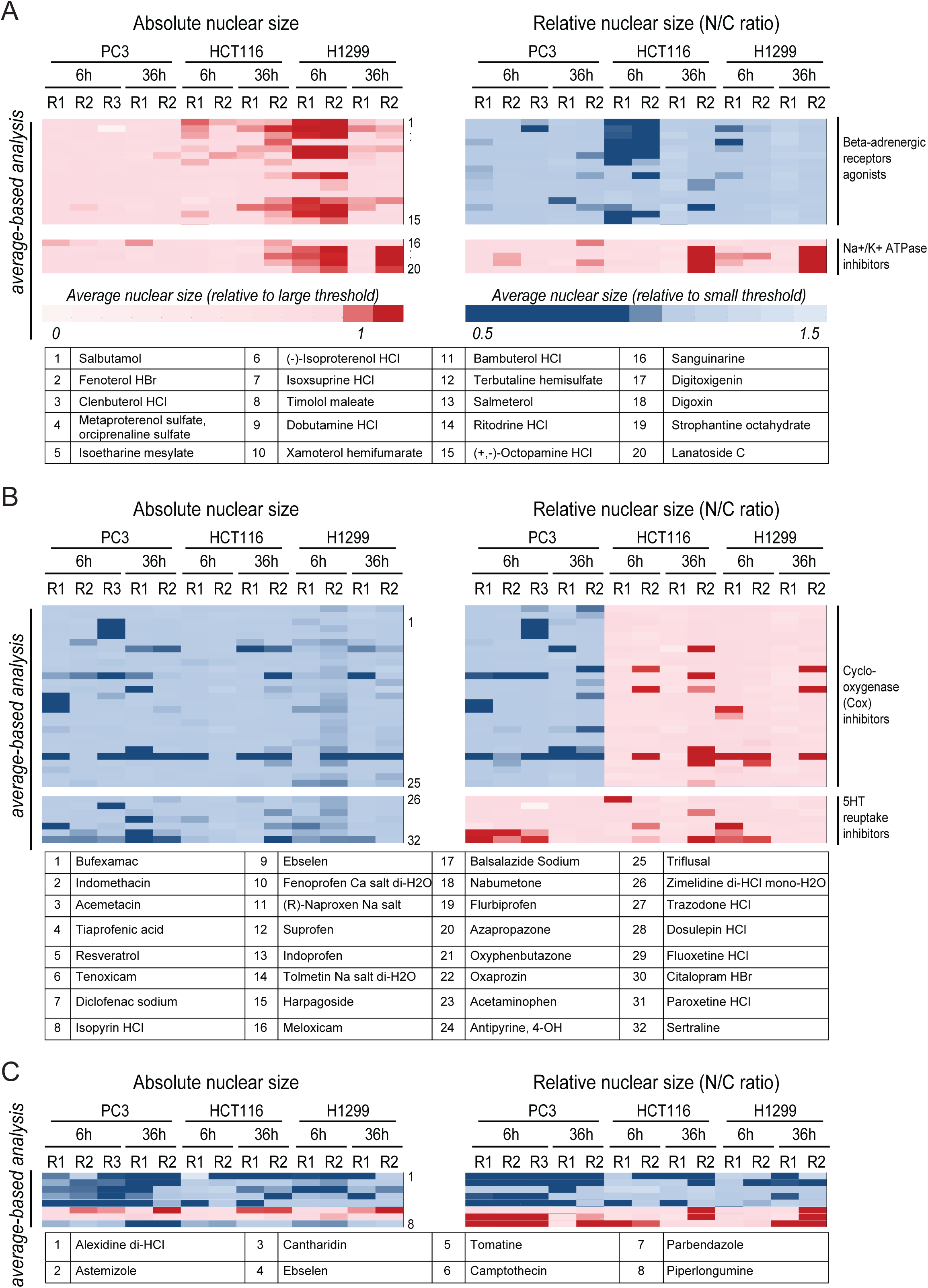
Functional groupings for NSR compounds segregated to particular tumor types. **a** Heat maps showing the mean nuclear size (left) and the mean N/C ratio (right) across tumor type/replicates (horizontal) treated with beta-adrenergic receptor agonists (top heat maps) or Na+/K+ ATPase inhibitors (bottom heat maps). Both heat maps show particular regions from Figures 3-4, magnified for better visualisation. Nuclear size and N/C ratio were normalised to the detection thresholds for large/small nucleus (see Methods), and colour-coded as indicated. Compounds are labeled from 1 to 20 and listed in the table below. Beta-adrenergic receptor agonists and Na+/K+ ATPase inhibitors correct nuclear size – N/C ratio in the right direction, the former increasing mean nuclear size in H1299 and decreasing N/C ratio in HCT116, while the latter increase both nuclear size and N/C ratio in H1299 cells. **b** Heat maps showing the mean nuclear size (left) and the mean N/C ratio (right) across tumor type/replicates (horizontal) treated with cyclooxygenase (Cox) inhibitors (top heat maps) or 5HT-reuptake inhibitors (bottom heat maps). Both heat maps show particular regions from Figures 3-4. Nuclear size and N/C ratio were normalised to the detection thresholds for large/small nucleus, and colour-coded as indicated. **c** Absolute versus relative nuclear size change profiles similarly extracted for eight other notable compounds. Compounds are listed in numerical order of appearance at the bottom of the panel.

We also identified NSR compounds that decreased nuclear size in PC3 prostate cancer cells including cyclo-oxygenase inhibitors and the 5-HT reuptake inhibitor class of anti-depressants (Fig. 5b). For the latter compounds (e.g., sertraline, paroxetine), the decrease in absolute nuclear size was sometimes accompanied by an increased N/C ratio after the early but not late time point, again consistent with potential homeostatic control mechanisms that act on the N/C ratio. Similar behaviour was observed for GABA receptor ligands/agonists on HCT116 colon cancer cells, whereby the drugs decreased absolute nuclear size but with an even greater effect on cell size, such that the N/C ratio was increased (Supplementary Figure S4b). The terpenoid natural product cantharidin, which has anti-viral and anti-proliferative effects, was a potential PC3 NSR that strongly decreased absolute nuclear size in PC3 cells while increasing the N/C ratio, both after 6 h and 36 h treatment (Figure 5c). In addition to the above illustrative examples, our nuclear size screens identified ∼50 other NSR compounds for the different cancer types (Table 1; Supplementary Figure S5). Thus, just as nuclear size changes are characteristic for each tumor type, many compounds that normalize these nuclear size defects are often specific to each cell type and exert effects on relative and/or absolute nuclear size metrics.

### NSRs have heterogeneous effects on cell viability

It is unclear whether nuclear size changes directly drive aggressive metastatic behaviour or if the alterations in nuclear size are a by-product of other aspects of tumor progression. To address this question, we selected 5 NSR compounds to test for properties relating to tumor growth and spread. The compounds selected covered a range of target specificities and directions of size changes (Table 2). Oxyphenbutazone, a non-steroid anti-inflammatory drug, which also depolymerises microtubules, decreased nuclear size in all three cell lines and at both 6 and 36 h time points, but specifically decreased the N/C ratio only in PC3 cells. Parbendazole, an anti-helminthic, increased relative nuclear size in all three cell lines but only at the 36 h time point. Digitoxigenin, a steroid lactone used to strengthen heartbeat that also has anti-cancer activity, dramatically increased nuclear size in H1299 cells but less so in HCT116 cells. Piperlongumine, an alkaloid natural product with claimed anti-cancer properties^32^, increased the N/C ratio in PC3 cells at 6h but decreased absolute nuclear size after 36 h and increased nuclear size in HCT116 and H1299 cells at 36 h. Finally, paroxetine, an anti-depressant that blocks serotonin uptake, was the most specific of the compounds tested as it caused a nuclear size decrease only in PC3 cells.

**Table 2.**
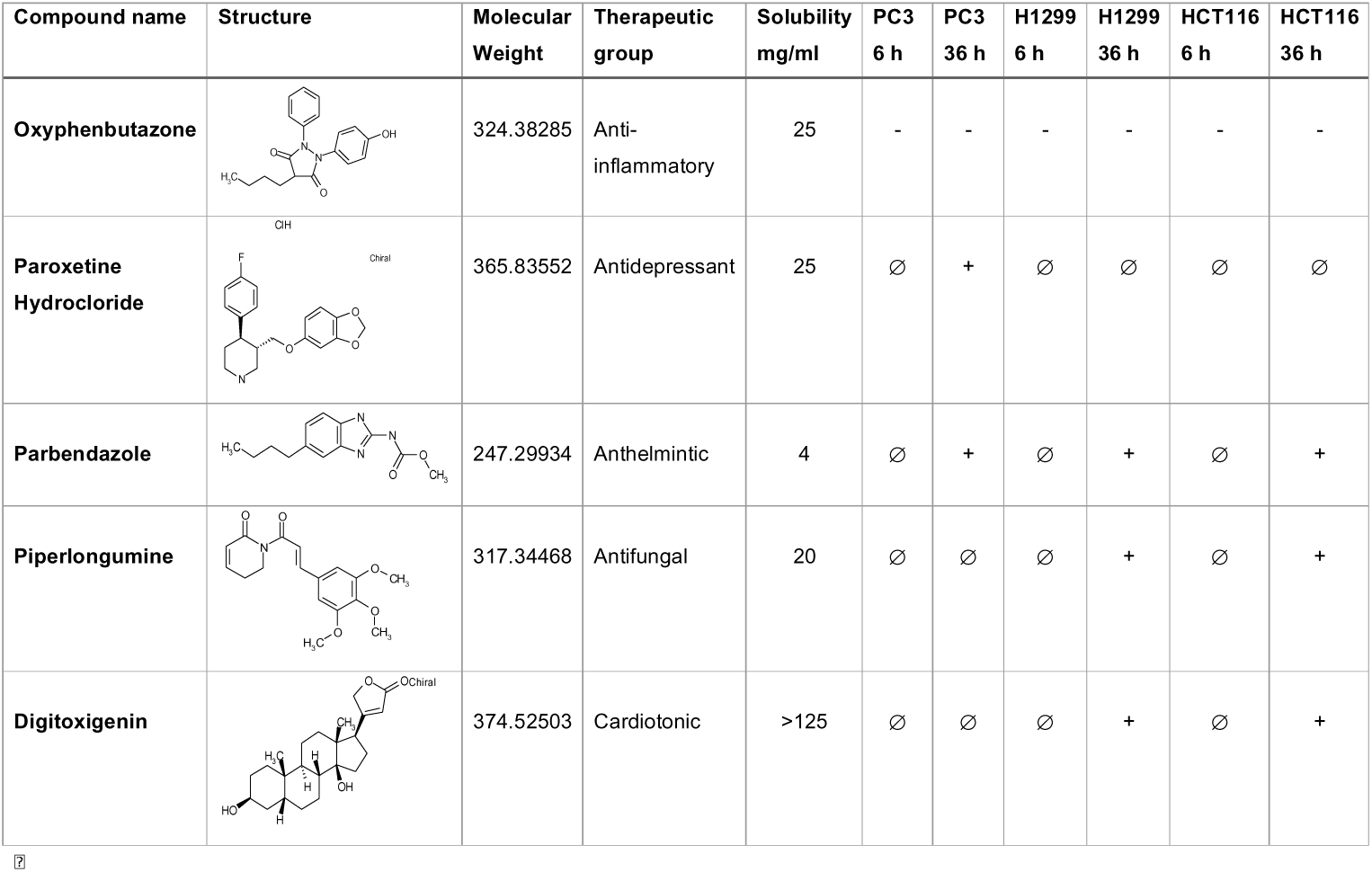
Compounds tested in detail with chemical structures.

| Compound name | Structure | Molecular Weight | Therapeutic group | Solubility mg/ml | PC3 6 h | PC3 36 h | H1299 6 h | H1299 36 h | HCT116 6 h | HCT116 36 h |
| --- | --- | --- | --- | --- | --- | --- | --- | --- | --- | --- |
| Oxyphenbutazone |  | 324.38285 | Anti-inflammatory | 25 | - | - | - | - | - | - |
| Paroxetine Hydrochloride |  | 365.83552 | Antidepressant | 25 | Ø | + | Ø | Ø | Ø | Ø |
| Parbendazole |  | 247.29934 | Anthelmintic | 4 | Ø | + | Ø | + | Ø | + |
| Piperlongumine |  | 317.34468 | Antifungal | 20 | Ø | Ø | Ø | + | Ø | + |
| Digitoxigenin |  | 374.52503 | Cardiotonic | >125 | Ø | Ø | Ø | + | Ø | + |

All five compounds reduced the viability and proliferative capacity of each cancer cell lines to different degrees, as assessed using an assay to quantify metabolically active cells in the population (Fig. 6a). Briefly, ∼5,000 cells were plated into each well of 96-well plates, drugs were added in serial dilutions and incubated for 36 h, then resazurine was added, which only reduces to fluorescent resorufine in metabolically active cells. Oxyphenbutazone decreased nuclear size in all three cancer lines and decreased cell viability similarly for PC3 cells and HCT116 cells, i.e., the two lines that exhibit a large nuclear size. In H1299 cells, which have a small nuclear size, viability decreased even more rapidly. In contrast, parbendazole, which increased nuclear size in all three lines had its strongest effect on viability of H1299 cells, i.e., in a context where it reversed the small nuclear size associated with tumor aggressiveness.

**Fig. 6.**
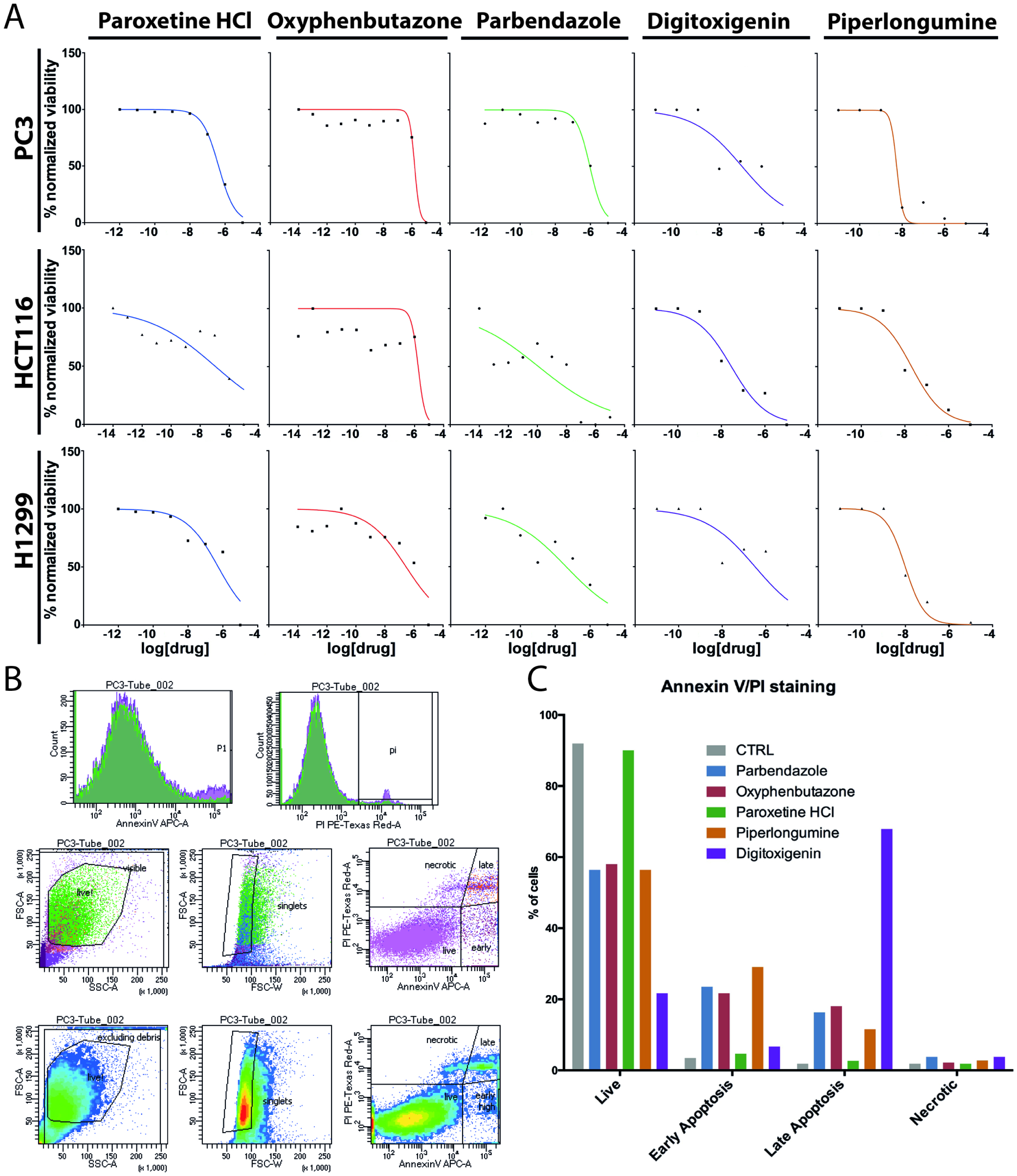
Proliferative and apoptotic effects induced by selected NSR compounds. **a** Paroxetine, oxyphenbutazone, parbendazole, digitoxigenin, and piperlongumine dose-response curves in PC3, HCT116 and H1299 cell lines. Cells were incubated on 96-well plates for 24 h with serial drug dilutions and DMSO controls. Resazurine cell viability solution was added to each well and fluorescence was assessed after 3 h of incubation and normalized to fluorescence signals of DMSO-treated control wells. Dose-response curves were obtained using least square fitting of the data points using Prism software. **b** Gating strategy for AnnexinV/Propidium Iodide (PI) FACS analysis. Forward scattering (FSC) vs Side scattering (SSC) plots (middle and bottom rows, left plots) were used to eliminate cell debris from the population, while distributions of AnnexinV and PI signals (top row plots) and FSC-Area vs FSC-width plots (middle and bottom rows, central plots) were used to remove cell doublets/triplets/groups. As a result, plotting individual cells according to their AnnexinV vs PI signals (middle and bottom rows, right plots) accurately identified populations of healthy (low AnnexinV, low PI), early apoptotic (finite AnnexinV, low PI), late apoptotic (finite AnnexinV, finite PI) and necrotic (low AnnexinV, finite PI) cells. **c** FACS results plotted as the percentage of the total cell population in each condition representing healthy, early apoptosis, late apoptosis, and necrosis at 36 h for drugs altering nuclear size analysed in this study. PC3 cells were treated with 10 μM of each compound for 36 h, refreshing the medium every day. Cells were as above and run through the FACS for detection of the two dyes.

To test if the concentrations that affected nuclear size might suffice to initiate apoptosis, we also performed Annexin V staining. As compounds that induce necrosis and consequent inflammation would have negative side effects if used therapeutically, we also stained cells with propidium iodide to distinguish necrotic cells. This co-staining protocol was applied to PC3 cells and resulted in 4 different cell populations: healthy cells with no staining, early-stage apoptotic cells positive only for Annexin V, late stage apoptotic cells positive for Annexin V and propidium iodide, and necrotic cells positive only for propidium iodide. Note that the first three populations are expected to contribute to resorufine fluorescence. The percentage of cells in each population was determined by fluorescence activated cell sorting (FACS). PC3 cells were incubated with drugs, refreshing the medium with drug every 24 h, and gating set to detect only singlets and intact cells (Fig. 6b). Plotting the percentage of cells in each category (Fig. 6c) revealed that digitoxigenin strongly triggered apoptosis with roughly 10% of cells in early apoptosis and ∼70% in late apoptosis. Parbendazole, oxyphenbutazone, and piperlongumine all yielded ∼40% cells in apoptosis, though with different percentages in early and late stages. Paroxetine, the only compound with a PC3-specific effect on nuclear size, induced neither apoptosis nor necrosis. From these results we concluded that there is no obvious correlation between effects on nuclear size and cell viability.

### NSRs inhibit cell migration and invasion

As nuclear size changes correlate with higher cancer grade and increased metastasis, we reasoned that drug-induced nuclear size corrections might affect cell migration or invasion. Size and scaling changes likely affect nucleoskeleton-cytoskeleton connections that in turn could affect cell migration, as previously indicated by slower cell migration with both knockout and overexpression of laminA, respectively in 2D and 3D assays^26, 27, 28, 33^. We tested a subset of the compounds identified in our screens for effects on cell migration in a scratch-wound healing assay (Fig. 7a-c). To be certain wound closure reflected cell migration and not cell division, cells were starved with 1% FBS containing medium 16 h before the scratch. Cells were treated with serial dilutions of compounds in four replicates immediately following the scratch and were recorded every 3 h for 48 h. Analysis of wound closure times against controls revealed that in PC3 cells oxyphenbutazone reduced migration in a dose-dependent manner at concentrations of 100 nM and higher. Parabendazole similarly inhibited migration but at higher concentrations of 1 µM or greater (Fig. 7a,b). Paroxetine did not have a quantifiable effect on PC3 cell migration in this assay system.

**Fig. 7.**
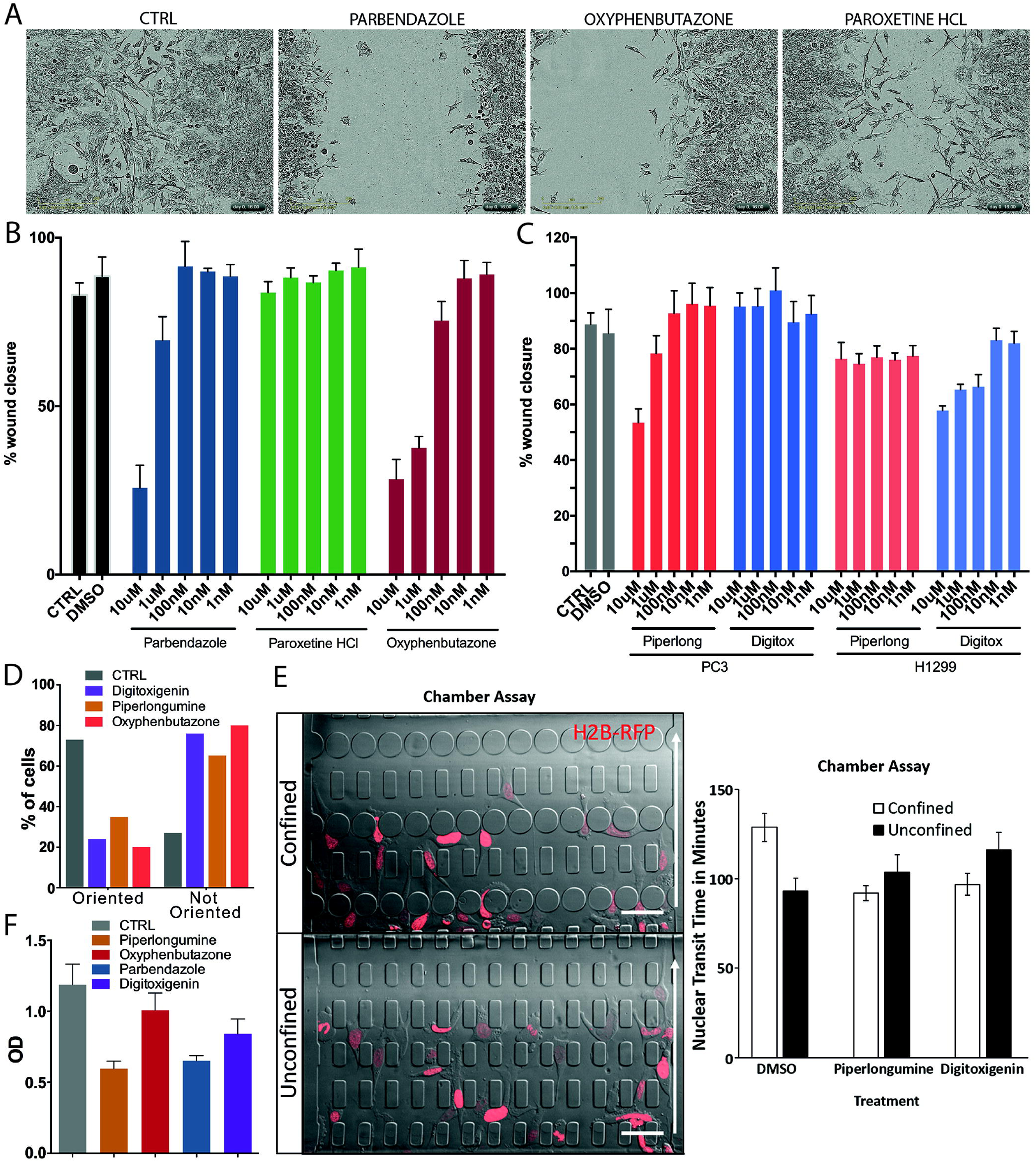
NSR compounds affecting nuclear size also affect cell migration. **a** Images of PC3 cells 24 h after scratch wound with 10 µM of indicated compounds. **b** Percentage of wound closure by PC3 cells after 24 h in the presence of serial dilutions of parbendazole, paroxetine or oxyphenbutazone quantified on an IncuCyte® system. **c** Percentage of wound closure by PC3 and H1299 cells in the presence of serial dilutions of piperlongumine or digitoxigenin similarly quantified. **d** Orientation assay. Percentage of cells from assays (B) and (C) after 12 h of wound closure in the presence of 10 µM of indicated compounds showing correctly oriented (left bars) or mis-oriented (right bars) centrioles. Centriole orientation was assessed using gamma tubulin staining. **e** Chamber assay. Representative images of cells migrating in the chamber assay used to measure rate of nuclear transit through confined (≤2 µm x 5 µm) or unconfined (15 µm x 5 µm) microfluidic constrictions. Arrow shows migration direction. Scale bar, 50 µm. Nuclear transit time is defined as the time required for cells to move their nucleus through an individual constriction. Error bars show SEM. Statistical significance between test groups was evaluated using a two-tailed t-test. The difference between any two test groups did not reach statistical significance (p<0.05). **f** Invasion assay. Cells were seeded in a Boyden chamber in absence of FBS and submerged on the outer chamber in presence of conditioned media and 10% FBS as chemoattractant. Cells were allowed to invade the membrane for 24 h prior to fixation and staining of the lower part of the membrane. Cells that had not invaded were wiped from the top of the membrane and the remaining cells that had invaded were lysed. The OD absorbance value for the lysed cells is plotted.

Considering that many compounds only affected nuclear size in certain cancer lines, we tested the other two compounds in both PC3 cells and H1299 cells. Piperlongumine also inhibited cell migration at 1 µM and higher in PC3 cells but had no effect on H1299 cells. Conversely, digitoxigenin inhibited cell migration of H1299 cells at 100 nM and higher, but exhibited no effect on PC3 cells (Fig. 7c). Unlike cell viability assays, the results of these migration assays were concordant with compound effects on nuclear size, i.e., compounds that rectified cancer-associated nuclear size changes (NSRs) also had anti-migration properties.

The effects in the wound healing assays could reflect reduced mobility or loss of directionality or both. To test for loss of cytoskeletal polarization, cells were stained for gamma tubulin to determine centriole orientation. After a scratch wound normal cells orient with the centriole facing the wound in order to direct inward migration and wound closure. Piperlongumine, digitoxigenin and paroxetine were tested in PC3 cells and each was found to cause a defect in centriole orientation in 65-85% of cells scored (Fig. 7d).

We considered the possibility that compound effects on nuclear size might reverse an advantage of the tumor cells to migrate through tight junctions. A weaker nucleoskeleton can enable squeezing through constrictions despite a larger nuclear size while a stiffer nucleoskeleton could enable smaller nuclei to move faster through constrictions^33, 34^. We tested the ability of the PC3 cells to migrate through 5×15 µm^2^ and 5×2 µm^2^ constrictions in the presence or absence of compounds. The small pore constrictions require substantial nuclear deformation, while the larger pores exceed the size of the nucleus and thus partly serve as controls for migratory ability in a 3D environment (Fig. 7e). The mean migration speed was slightly slower for piperlongumine-treated cells through the 5×15 µm^2^ constrictions (79 min) versus DMSO solvent control (91 min). Similarly, digitoxigenin-treated cells exhibited a slightly slower mean migration speed than controls (73 min versus 82 min) through the smaller 5×2 µm^2^ constrictions. However, in both cases the difference did not reach statistical significance. As cells do not need nucleo-cytoskeletal connections to squeeze through the constrictions and the centriole was not oriented towards the scratch in the wound healing assays, the lack of statistical significance suggests that the slower migration in the scratch wound assays may arise from defects in 2D cell polarization. As cells can use different migration modes in 2D wound healing assays versus 3D chamber migration assays^33, 34, 35^, these results are consistent with the effects of the compounds in wound healing assays but suggest that the effects have more to do with reduced mobility and directionality than with a change in mechanical stiffness.

A transwell migration assay was performed (Fig. 7f) to investigate compound effects on PC3 invasion. Cells were seeded on the inside Boyden chamber in absence of FBS on a 3 µm pore membrane coated with ECM proteins and submerged on the outside Boyden chamber in conditioned media as chemoattractant. Cells were allowed to migrate through the membrane for 24 h in the presence of compounds at the 10µM concentration used in the size screen, and the relative number of cells migrating was quantified with a colorimetric assay. Cells treated with piperlongumine and parbendazole had ∼50% fewer cells migrate through the membrane compared to DMSO-treated controls, where oxyphenbutazone and digitoxigenin reduced the invasion capability by 15% and 30% respectively. Thus, NSR compounds reduced both cell migration and invasion in tissue culture assays.

## Discussion

We postulated that nuclear size changes that correlate with increased metastasis might contribute directly to metastasis through changes in cellular mechanics and/or migration. Screening for compounds that alter nuclear size in 3 distinct cancer types revealed different sets of compounds affecting each and most of the compounds tested inhibited cell migration. Many of the FDA/EMA-approved compounds that we show have cell line-specific NSR activity in reversing the direction of cancer size changes have not been previously tested for cancer therapies for these tumor types. Our data indicated that DNA intercalants and microtubule polymerisation inhibitors, commonly used in anti-cancer treatments, affect nuclear size regulation non-specifically in a broad range of tissues, which may play a role in their extensive side effects. Such systemic toxic side effects are the principle problem with current chemotherapy regimens. In contrast, our tumor-type/tissue-specific targeting compounds may improve tissue-specificity of chemotherapy regimens, by targeting tumor spread while reducing systemic toxicity.

The tumor-type/tissue-specificity observed here for different NSRs has important ramifications for understanding mechanisms of scaling/size regulation, tissue function, and tumor metastasis. The evolutionary conservation of cell and organelle scaling indicates its general importance, yet little is known about scaling mechanisms. The finding of compounds that affect average nuclear size differently from N/C ratios should open a new avenue of research into scaling mechanisms. At the same time, our data indicates that in addition to evolutionarily conserved pathways (reviewed in^24^), nuclear/cell scaling and nuclear size regulation have tissue-specific components. It is tempting to speculate that, as higher organisms developed more specialised tissues, the unique connections of the NE for different tissue functions became entwined with scaling regulation. For example, an ovary needs a different scaling regulation to support nutrient storage while colonic epithelia have polarized cytoskeleton connections to the nucleus. These mechanisms could be hijacked in tumorigenesis to achieve observed tissue-specific nuclear size changes. Such differences in mechanism and function could impact on whether or not the size change confers an advantage to the cancer in a particular cell type.

Many compounds not only differed in their tissue-specific effects on nuclear size but also in their effects on absolute versus relative nuclear size. We speculate that disruption of nuclear scaling (i.e., N/C ratio) might be a fundamental aspect of metastasis such that reversing the scaling defect might yield a less aggressive tumor. However, even if the scaling change is not fundamental to tumorigenesis, restoring normal nuclear size might prevent metastatic spread if the initial size change altered nuclear plasticity to facilitate squeezing through constrictions (Fig. 8a) or increased tumor cell motility (Fig. 8b).

**Fig. 8.**
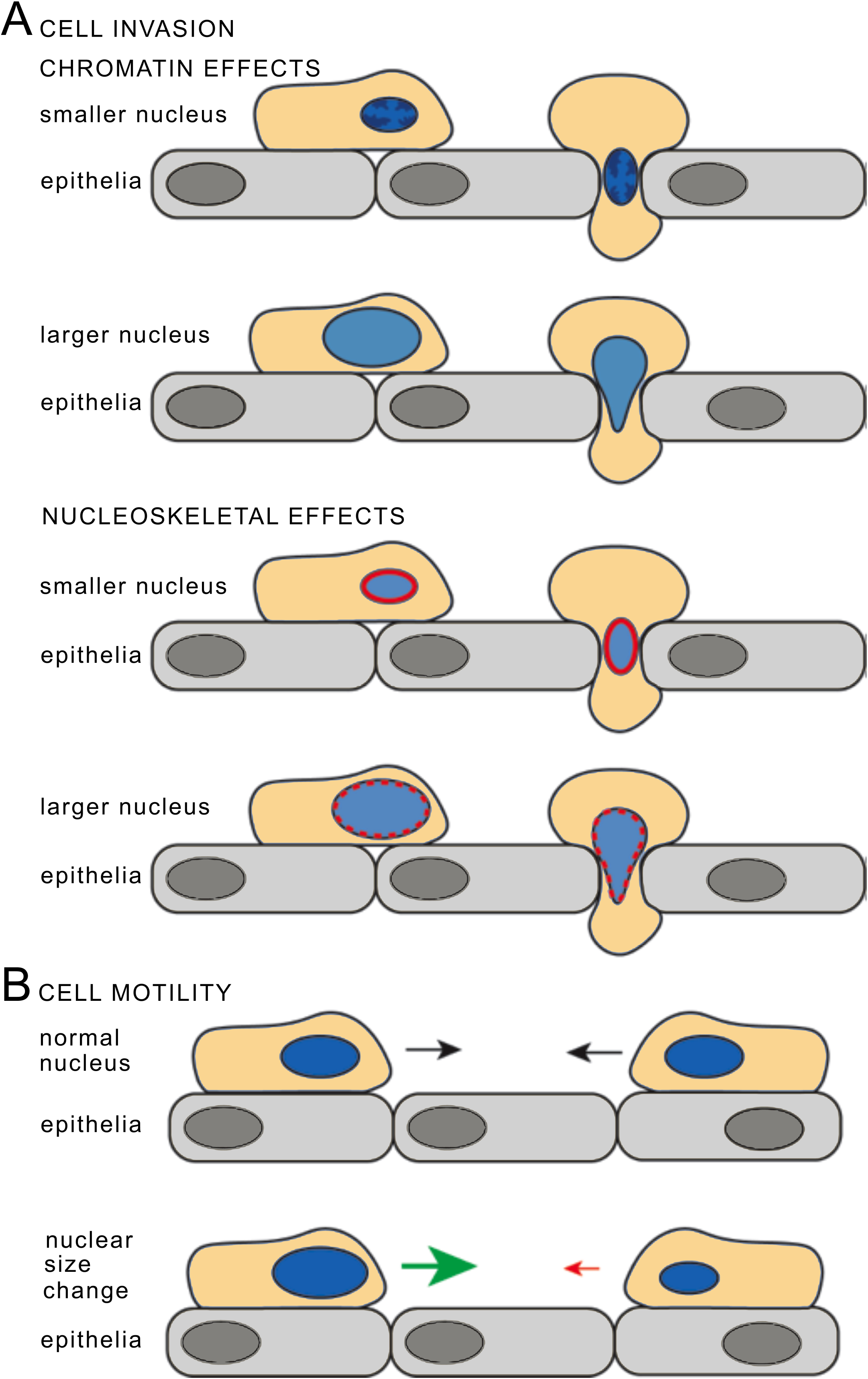
Model for how nuclear size changes may affect cancer cell invasiveness. **a** Chromatin effects on nuclear size and stiffness (top). For cell invasion through an epithelial layer, condensation of chromatin in a smaller nucleus could allow facile transit through a tight junction while looser chromatin in a larger nucleus could make the nucleus more malleable and also facilitate transit through a tight junction. Nucleoskeletal effects on nuclear size and stiffness (bottom). Reduced lamin A production might result in a smaller nucleus that could squeeze through tight junctions. Conversely scaling defects that cause a larger nucleus with lower lamin density would be more malleable to facilitate tight junction transit. **b** Cell motility may also be affected by nuclear size. Altered connections between the nucleoskeleton and cytoplasmic filaments in cells with larger or smaller nuclear size may influence cell mobility.

The nucleus is the largest and sturdiest organelle and thus the greatest impediment to cell migration through tight constrictions, such as when a metastatic cell migrates into other tissues. Making the nucleus smaller could promote metastasis by facilitating cell squeezing through cell-cell junctions while a larger nucleus —without an increase in ploidy so that the DNA is less compact— may more readily deform to squeeze through cell-cell junctions (Fig. 8a). Moreover, simply changing the nature or density of connections between the nucleus and cytoskeleton could alter cell migration. Indeed, the actin networks at the leading edge connect all the way back to the nucleus and actin networks exert profound effects on nuclear stiffness and cell migration, as well as tumorigenesis itself^27, 36, 37^. Thus, either reversing specific nuclear size changes in a particular cancer type or further exaggerating the nuclear size change could re-impose fitness bottlenecks on tumour evolution.

Precisely how many NSR compounds affect nuclear size may diverge from established or putative mechanisms of action. Although oxyphenbutazone is commonly used as an anti-inflammatory, it caused cells to round up, even at 6 h when apoptotic effects were negligible. This may reflect a second function inhibiting Wnt/β-catenin as reported in hepatocellular carcinoma model systems^38^. Likewise, digitoxigenin induces cytotoxic effects on non-small cell lung cancer cells through a second function inhibiting the Na^+^/K^+^ ATPase activity. Digitoxigenin is an effective chemotherapeutic against metastatic uveal melanoma^39, 40^ and its derivative, amantadig, is effective in inhibiting hormone-refractory prostate cancer cells^41^, but the mechanism by which it acts is not clear and may even differ in different cancer types. Piperlongumine has been used to treat breast cancer in xenograft models, causing a reduction of tumor volume and metastasis^42, 43^ and this activity is thought to function through ROS-mediated apoptosis^32, 43, 44, 45^. Piperlongumine also has also been reported to inhibit glycolysis and FOXO3A, and to have senolytic and antimicrobial activity^46, 47, 48, 49^. How these mechanisms relate to piperlongumine effects on nuclear size remain to be determined.

The other compounds we tested for migration effects have not been used to our knowledge as anti-cancer agents, nor can their known targets readily explain their effects on nuclear size regulation. Paroxetine is an antidepressant of the selective serotonin reuptake inhibitor (SSRI) class and is used to treat depressive and obsessive-compulsive disorders. Paroxetine interacts with cytochrome P450 complex enzymes^50^ and promotes mitochondrial-induced apoptosis in astrocytes^51^. Parbendazole is a benzimidazole derivative showing broad-spectrum anthelminthic activity and is also a potent microtubule assembly inhibitor^52, 53^. Parbendazole induced apoptosis in nearly 40% of PC3 cells and inhibited cell migration by nearly 75% in wound healing assays and by 50% in invasion assays. Thus, even though this compound is from a group of drugs with anti-helminthic activity, it may hold promise as an anti-cancer therapeutic.

Other anti-helminthics affected nuclear size in our study as well as several other compound classes not further investigated. Thus, several compound classes not previously used in cancer treatments might be repurposed to treat particular cancer types. For some compound classes this has already been achieved. For example, BAAs were NSRs for colonic adenocarcinoma and small cell lung cancer cells. Interestingly, BAAs suppress the epithelial to mesenchymal transition of bronchial epithelial cells and have been used to treat lung cancer^54, 55^. BAAs also inhibit cancer pathways for triple-negative breast cancer and gliomas^56, 57^, and the agonist salbutamol reduces migration and invasion of breast cancer cells^58^. Tissue-specific context for BAAs may be critically important, however; for example, beta-adrenergic receptor signaling promotes angiogenesis of gastric tumors^59^. Serotonin reuptake inhibitors are another NSR class that has recently been repurposed to treat glioblastoma^60^ and shown promise in model systems against hepatocellular carcinoma^61, 62^.

This study provides proof-of-principle to the hypothesis that targeting tissue/cancer type nuclear size changes can affect cell migration with the potential to reduce metastasis. Accordingly, similar screens should be engaged for other cancer/tissue types and the compounds identified tested in animal models. The new compounds and compound classes identified in this study will likely enhance anti-cancer therapies by increasing tissue-specificity, reducing toxicity, and directly targeting metastasis. Due to their tissue-specificity, combining them with current chemotherapy regimens should allow for reduction of some of the compounds that target all cell types and thus yield systemic toxicity. Most critically, the tendency of compounds affecting nuclear size to also reduce cell migration and invasion suggests that their inclusion in treatment regimens could reduce the metastasis that is usually the primary cause of mortality and thus improve long-term survival.

## Methods

### Cell lines

PC3, HCT116, and H1299 cells were stably transfected to express H2B-mRFP in the pEGFP-N2 clontech vector and maintained under G418 selection (500_µg/mL) for general culture. Selection was removed for 1 passage prior to plating for experiments to avoid drug interference effects.

### Cell culture and manipulation

The Prestwick compound library was contained in fourteen 96-wells plates, each of which consists of 80 drugs mostly approved by the FDA and EMA agencies (1 mM concentration in DMSO) and 16 in-plate DMSO controls, for a total of 1,120 compounds. For each screen, ∼5,000 H2B-mRFP tagged cells were plated on each well of fourteen Greiner Screenstar 96-wells glass-bottom imaging plates and left overnight in 99 µL RPMI medium for adhesion and growth prior to compound treatment. Then compounds were added by pipetting directly 1 µL of 1mM DMSO stock directly into the plate wells (final compound concentration: 10 µM), and cells were incubated 6 h or 36 h before preparation for imaging. Cell densities prior to compound treatment were adjusted to account for cell doubling during the 36 h long-term treatment in order to minimize differences in cell densities at the time of imaging. Cell were fixed in 3.7% formaldehyde fixation solution (EMD FX0410-5 formaldehyde solution) for 15 min at room temperature, washed with PBS, then incubated 30 min µl HCS CellMask Deep Red 0.5X (Molecular Probes) for cytosol staining. Stained fixed cells were washed again and stored in PBS at 4°C. Plates were wrapped in parafilm to prevent PBS evaporation and in aluminium foil to prevent fluorochome damage by ambient light, for a maximum of one week before imaging.

### Imaging data acquisition

H2B-mRFP tagged cells were imaged using an Opera^TM^ High Content Screening instrument (PerkinElmer), equipped with a 20x air objective (LUCPLFLN, NA=0.45). H2B-mRFP (nucleus) and CellMask DeepRed (cytosol) fluorochromes were excited sequentially by 561 nm and 640 nm lasers respectively, and emitted fluorescence was acquired by Peltier-cooled CCD cameras under two different channels with bandpass detection filters centred at 600 nm (600/40) and 690 nm (690/70) with respective exposure times of typically 320 ms and 200 ms. Bleed-through of the cytosolic channel into the nuclear channel was negligible. For each well, 20 Fields of View (FOV) were acquired at the same positions for all wells and plates, avoiding the edge of the well where cells accumulate due to capillary action. This allowed quantification of nuclear and cellular size for several hundreds of cells for each condition/replicates. For technical reasons, the two replicates of the 6 h treatment time point for the H1299 cell line were acquired on an equivalent Opera Phenix imaging system (PerkinElmer). Quantitative analysis of the data was carried out using the metrics and analysis methods detailed below. An adapted Acapella^®^ (PerkinElmer) software script was used to automatically mask the cytoplasm and nucleus of individual cells, and exported single cell results to a .txt file for subsequent analysis in MATLAB (The Mathworks, see below).

### Data processing and quantitative analysis

#### i) Data filtering and computation of nuclear size metrics

Individual cells and nuclei within each FOV were masked using the automated threshold-based detection algorithm on the Opera platform (Acapella^®^ scripting environment). Intensity threshold filters, size filters, and morphological filters (which threshold on cell/nuclei size, nuclear roundness, width to length ratio, distance between nuclei) were used to filter out detection artefacts, multiple detection of single cell/nuclei or unique detection of cell clusters. Cell and nuclear sizes were respectively defined as the areas (in pixels) of the masked regions in the focal plane, based on respectively the CellMask DeepRed and H2B-mRFP signals. Individual cell output data, including cell and nuclear sizes, and average signal intensities in the two compartments were saved as text files and loaded in MATLAB for statistical analysis (Fig. 1, see below). Identical detection and filtering parameters were used in Acapella^®^ to identify cells and nuclei under all conditions to homogenize data analysis and prevent post-acquisition processing biases. For conditions that yielded higher cell densities at the time of imaging, the above filtering step was sometimes insufficient to remove clusters of cells so cell size outliers were eliminated from all datasets. For each condition separately, the entire cell size distribution was analysed and we extracted the 25%, 50% and 75% size quartiles (Q1, Q2 and Q3 respectively). Following standard outlier removal procedures, we filtered out detection areas with a size larger than the outlier threshold Q2+6*(Q3-Q2) (i.e., likely cell clusters), and the detections smaller than Q2-6*(Q2-Q1) (i.e., imaging artefacts or cell debris). All DMSO control wells for each plate were processed together in this step.

For each detected cell that passed through data filtering steps, we computed two metrics: the absolute nuclear size N, and the relative nuclear size, or N/C ratio, defined as the ratio of the absolute nuclear size to the absolute cell size (i.e., the area of the CellMask DeepRed signal focal cross-section in pixels).

#### ii) Statistical analysis of distributions - average-based and outlier-based strategies

Compounds that altered absolute or relative (N/C) nuclear size metrics were identified using two complementary strategies for each metric.

##### Average-based strategy

first, for each plate/replicate, the average value of the metric across all detected cells was determined for each compound-treated well and for all in-plate DMSO control wells. Hits were selected as compounds with an average metric value that stands out from the distribution of average metric values across the library. Specifically, we computed the 25%, 50% and 75% percentiles of the distribution of well-averaged metric values across the library for each cell line/replicate separately (respectively denoted Q1, Q2 and Q3). Then low and high outlier thresholds were computed following standard procedures, respectively as *mean_low_* = *Q*2 – *s* * (*Q*2 – *Q*1) and *mean_high_* = *Q*2 + *s* * (*Q*3 – *Q*2). In standard procedures, the parameter *s* is generally chosen as *s* = 6 for hard hit selection and *s* = 3 for soft hit selection. We used the former for the absolute nuclear size metric and the latter for the N/C ratio metric. N/C ratio cell-to-cell variability exceeded absolute nuclear size cell-to-cell variability, likely because the N/C ratio originates from 2 different measurements (nuclear size and cell size), both of which are associated with measurement uncertainty and biological variability. Hence, in order not to reduce the hit discovery potential of the N/C metric, we used slightly less stringent criteria for hit selection with this metric. This may explain, in part, the increased sensitivity of the N/C ratio relative to nuclear size in identifying active compounds. These pairs of thresholds are represented as dashed red lines on Supplementary Figure S1B, top row and bottom row for nuclear size and N/C metrics, respectively. Finally, compounds that reduce (respectively increase) the mean value of the metric beyond non-specific effects are compounds that stand out of the distribution of mean metric values across the Prestwick collection by respectively reducing the metric below / increasing the metric above the low and high thresholds *mean_low_* and *mean_high_*. Such compounds were designated as hits for the corresponding metric and the *average-based* analysis strategy.

##### Outlier-based strategy

For each plate, we computed the 25%, 50% and 75% percentiles characterizing the distribution of metric values across the population of individual DMSO-treated control cells only (i.e., 16 wells per plate). We used these percentiles (Q1, Q2, Q3) to compute low and high (soft) outlier thresholds as defined above *DMSO_lowThres_* = *Q*2 – 3 *(*Q*2 – *Q*1) and *DMSO_highThres_* = *Q*2 + 3 *(*Q*3 – *Q*2). Cells with values below the low threshold were designated as having an abnormally small metric; likewise, the high threshold represents the metric value above which a cell was designated as having an abnormally large metric. We next used these thresholds to compute, for all DMSO control and compound wells, the fractions of cells showing abnormally small and large metric values (relative to the in-plate DMSO distribution), i.e. such that *Metric* < *DMSO_lowThres_* or *Metric* < *DMSO_highThres_* respectively (Fig. 1B, central plot, red-shaded areas). These fractions were typically 0-10% for DMSO controls and cells treated with compounds showing no specific effect on nuclear size. These numbers reflected the biological and measurement-related cell-to-cell variabilities in our dataset. However, these fractions could reach 20-30% of the cells or even more for the strongest compounds (e.g. oxyphenbutazone, Fig. 1B right plot). Thus, compounds that deregulate the metric beyond the expected variability in our entire dataset are compounds that enrich one or both of those fractions of outlier cells above this typical level defined by DMSO controls and other compounds. To detect these compounds, we computed the entire distributions of each of these fractions of outlier cells separately across the entire compound collection, computed the 25%, 50% and 75% percentiles that characterize the distributions of those fractions of outlier cells with abnormally small/large metric value across the library (Q1_small/large_, Q2_small/large_ and Q3_small/large_), and defined hit detection thresholds as *Frac_small_* > *Q*2*_small_* + 6 * (*Q*3*_small_* – *Q*2*_small_*) and *Frac_large_* > *Q*2*_large_* + 6 * (*Q*3*_large_* – *Q*2*_large_*). We note that the qualitatively different nature of the analyses and the corresponding thresholds used could explain differences in hit counts between average and outlier analyses.

#### iii) Hit confirmation

Hits were confirmed by computing direct compound-vs-DMSO Wilcoxon rank tests, where the multiple well-averaged values of the metric across all DMSO and compound replicates were used to perform the tests. Reported p-values account for variability between plates in well-averaged metric values from both DMSO and compound-treatments. In some rare instances, this variability precluded the identification of some weaker compounds.

### Pilot screen

To estimate the accuracy of these analysis strategies and assess screen reproducibility, we ran a pilot screen on PC3 cells at the 6 h time point in triplicate (Table S1 and Supplementary Figure S1). For each hit, obtained using absolute or relative (N/C ratio) nuclear size metrics (rows 1-2 and 3-4 respectively), the distributions of mean metric values (rows 1-3) and fraction of outlier cells (rows 2-4) across the 3 replicates (thus, 3 values per metric-analysis strategy pair) were compared to the similar distributions from all DMSO controls of the collection using a Wilcoxon rank test. These tests yielded for every hit a p-value score that is represented as scatter plots on Supplementary Figure S1A. Among hits replicated exactly once, twice or three times, the fraction of hits showing significant (p<0.05) p-values are shown on Table S1. As a result, ∼85-100% of the hits that replicated twice and three times had low p-values irrespective of the metric/analysis strategy. A large fraction of hits that scored only once out of three replicates still showed statistical significance upon direct comparison with DMSO controls, in particular for the nuclear size metric analysed with the average-based strategy. This reflected scores slightly below hit detection thresholds, as can be viewed on scatter plots showing the raw data across replicates for three plates (240 compounds, Supplementary Figure S1B). Of note, some compounds that never scored above detection thresholds nevertheless did show statistically significant altered nuclear size. However, these compounds were not analysed further. The rather low false discovery rate <20% for hits determined using *average-based* analysis of absolute nuclear size prompted us to downsample larger screens to duplicate screens, and to use this metric and analysis method as the default hit discovery methodology. Relative nuclear size metric (as quantified by the N/C ratio) and *outlier-based* analysis were used as indicated in the main text to provide additional contrasts to emphasize the phenotypic specificity of particular compound classes.

### Apoptosis assay

Cells were incubated with each compound at a concentration of 10 μm standard 6-well tissue culture plates. Approximately 10^6^ cells were counted, washed with ice plates. Approximately 10^6^ cells were counted, washed with ice cold PBS and stained with 5 μL of the Annexin V apoptosis marker conjugate with the 647-Alexafluor chromophore (Thermo Fisher Scientific) and 5 μL of 5 μg/mL Propidium Iodide (Biotium) for cell death detection, in 10 mM HEPES, 140 mM NaCl, and 2.5 mM CaCl_2_, pH 7.4 for 15 min before FACS analysis. The 647 nm Annexin signal was used for detection of early stages of apoptosis as cells are not permeable to propidium iodide at this stage. Late stage apoptosis was detected by presence of both of the markers and necrotic cells by the presence of propidium iodide signal only.

### Viability assay

For viability analysis a resazurine-based assay was performed. This assay allows quantification of live cells due to reduction of the resazurine (a.k.a. Alamar Blue) to resorufine, a red and highly fluorescent compound in metabolically active cells. The detection of either the absorbance or fluorescence of the coupled resazurine/resorufine allows the determination of live cell numbers compared with controls. The cells were seeded onto PE96 plates (PerkinElmer) at a concentration of 5,000 cells/well in a final volume of 100 μL and allowed to recover/grow for 24 h before compound addition. Media was then replaced with media containing six serial dilutions of each compound and cells were allowed to grow for either 6 or 36 h. 10 μL Alamar blue (Thermo Fisher, 10x stock) reagent was added on each single well and allowed to react for 3 h prior to taking an absorbance reading at 530 nm or a fluorescence reading with excitation at 560 nm (substrate excitation 530-570 nm) and emission at 590 nm (substrate emission 580-590 nm) on a microplate reader (JASCO V-550). The resorufine extinction coefficient E^mM^ (572 nm) is 73 in pH 8.0. The concentration of reagent in the Alamar blue solution is proprietary, but it must be much higher than these standard drug concentrations to work in this widely used assay such that compound fluorescence in this range would be unlikely to alter measurement values.

### Wound healing assay

Approximately 25,000 cells per well were seeded on a Sartorius 96-well LockView plate (Sartorius) the night prior to wound formation. The cell monolayer was scratched with the IncuCyte® WoundMaker, that simultaneously makes equivalently-sized scratch wounds in the monolayer in all wells, and medium replaced with compounds containing medium supplemented with 1% FBS to induce cell migration and reduce cell proliferation. Plates were placed in the IncuCyte® incubator and imaged in bright field every 3 h for 48 h. Images shown in Figure 7a were generated using the IncuCyte ZOOM while subsequent quantification was done using the IncuCyte® S3. Analysis of the wound closure time were performed with an automated script provided by Sartorius to determine the percentage of wound closure.

### Microfluidic device migration assay

PC3 cells were seeded into a previously described microfluidic device for studying nuclear transit during confined migration^63^. The device consists of migration channels with a fixed height of 5 µm and constrictions of 1 to 2 µm or 15 µm in width. Devices were assembled as previously described and coated with a solution of 5 µg/mL of fibronectin 24 h prior to experiments^64^. 30,000 cells were seeded into each device 6 h prior to imaging and were treated with either piperlonguine, digitoxigenin, or DMSO control. Imaging was performed on a Zeiss LSM700 laser scanning confocal microscope with a 20x air objective. Cells and devices were imaged at 10 min intervals in a temperature controlled stage (37°C) for 14 h. The time required for cells to migrate through an individual constriction was quantified using a previously described MATLAB script for measuring nuclear transit through these microfluidic devices^65^.

### Invasion assay

The QEM Endothelial cell invasion assay was carried on in accordance with manufacture’s recommendations (ECM210, Millipore). Briefly, cells were seeded at a concentration of 10^5^ on the inner part of a Boyden chamber in absence of FBS and submerged on the outer chamber in presence of conditioned media and 10% FBS as chemoattractant. Cells were allowed to invade the membrane for 24 h prior fixation and staining of the lower part of the membrane. Non-invading cells were removed with a cotton stab form the top of the membrane and cells lysed. Absorbance values of the cell lysate were analysed on a microplate reader (JASCO V-550) with a 540 nm wavelength.

## Data availability

Data generated or analysed during the current study are included in the published manuscript and its supplementary information files.

### Online Content

Methods, Supplementary Figures S1-5, Supplementary Tables S1-5 with statistics and detailed listings of all compounds identified by average-based and outlier-based approaches.

## Acknowledgments

We thank Edward Jarman and Bernard Hormann for initiating pilot studies as part of their undergraduate Honours and MSc diplomas, respectively. We also thank the McGill University core facility for allowing us to perform some imaging on their Opera Phenix system and the Institute for Research in Immunology and Cancer, University of Montreal, High Throughput Core Facility for general support. This work was principally supported by Wellcome grants 095209 (E.C.S.) and 085178 (M.T.), the European Research Council SCG-233457 (M.T.), the Canadian Institutes of Health Research FDN-167277 (M.T.) a Canada Research Chair in Systems and Synthetic Biology (M.T.) and Medical Research Council MR/R018073/1 to ECS. It was also supported by Wellcome Centre for Cell Biology 092076, the National Institutes of Health R01HL082792 and U54CA210184 (J.L.), the Department of Defense Breast Cancer Research Program Breakthrough Award BC150580 (J.L.), and the National Science Foundation CAREER Award CBET-1254846 (J.L.). A.R. was supported by a Darwin Trust Studentship and J.T.K. from the Knight@KIC 2018-2019 Graduate Fellowship. This work was performed in part at the Cornell NanoScale Science & Technology Facility (CNF), a member of the National Nanotechnology Coordinated Infrastructure (NNCI), which is supported by National Science Foundation (Grant NNCI-1542081).

## Author Contributions

Conceived/ designed experiments: AR, ST, MT, ECS. Performed experiments: AR, ST, NTP, NZ, JTK, DB, JC, SZ. Provided analysis tools and scientific input: ST, JW, NTP, PC, JL, MA, NOC, VGB, MT. Wrote the manuscript: ST, AR, MT, ECS. All authors contributed to editing the manuscript.

## References

1. Beale LS. Examination of sputum from a case of cancer of the pharynx and the adjacent parts. Arch Med Lond 2, 44 (1860).

2. de Las Heras JI, Batrakou DG, Schirmer EC. Cancer biology and the nuclear envelope: A convoluted relationship. Semin Cancer Biol 23, 125–137 (2013).

3. de Las Heras JI, Schirmer EC. The nuclear envelope and cancer: a diagnostic perspective and historical overview. Advances in experimental medicine and biology 773, 5–26 (2014).

4. Zink D, Fischer AH, Nickerson JA. Nuclear structure in cancer cells. Nat Rev Cancer 4, 677–687 (2004).

5. de Andrea CE, Petrilli AS, Jesus-Garcia R, Bleggi-Torres LF, Alves MT. Large and round tumor nuclei in osteosarcoma: good clinical outcome. Int J Clin Exp Pathol 4, 169–174 (2011).

6. Ladekarl M, Boek-Hansen T, Henrik-Nielsen R, Mouritzen C, Henriques U, Sorensen FB. Objective malignancy grading of squamous cell carcinoma of the lung. Stereologic estimates of mean nuclear size are of prognostic value, independent of clinical stage of disease. Cancer 76, 797–802 (1995).

7. Abdalla FB, Markus R, Buhmeida A, Boder J, Syrjanen K, Collan Y. Estrogen receptor, progesterone receptor, and nuclear size features in female breast cancer in Libya: correlation with clinical features and survival. Anticancer research 32, 3485–3493 (2012).

8. Nandakumar V, et al. Isotropic 3D nuclear morphometry of normal, fibrocystic and malignant breast epithelial cells reveals new structural alterations. PLoS One 7, e29230 (2012).

9. Rashid F, Ul Haque A. Frequencies of different nuclear morphological features in prostate adenocarcinoma. Annals of diagnostic pathology 15, 414–421 (2011).

10. Tan PH, Goh BB, Chiang G, Bay BH. Correlation of nuclear morphometry with pathologic parameters in ductal carcinoma in situ of the breast. Modern pathology : an official journal of the United States and Canadian Academy of Pathology, Inc 14, 937–941 (2001).

11. Chen L. Cytology: diagnostic principles and clinical correlates. The American Journal of Surgical Pathology 34, 286 (2010).

12. Cavalier-Smith T. Economy, speed and size matter: evolutionary forces driving nuclear genome miniaturization and expansion. Ann Bot 95, 147–175 (2005).

13. Edens LJ, White KH, Jevtic P, Li X, Levy DL. Nuclear size regulation: from single cells to development and disease. Trends Cell Biol 23, 151–159 (2013).

14. Fidorra J, Mielke T, Booz J, Feinendegen LE. Cellular and nuclear volume of human cells during the cell cycle. Radiat Environ Biophys 19, 205–214 (1981).

15. Jorgensen P, Edgington NP, Schneider BL, Rupes I, Tyers M, Futcher B. The size of the nucleus increases as yeast cells grow. Mol Biol Cell 18, 3523–3532 (2007).

16. Neumann FR, Nurse P. Nuclear size control in fission yeast. J Cell Biol 179, 593–600 (2007).

17. Jevtic P, Edens LJ, Li X, Nguyen T, Chen P, Levy DL. Concentration-dependent Effects of Nuclear Lamins on Nuclear Size in Xenopus and Mammalian Cells. J Biol Chem 290, 27557–27571 (2015).

18. Lu W, et al. Nesprin interchain associations control nuclear size. Cell Mol Life Sci 69, 3493–3509 (2012).

19. Kume K, Cantwell H, Burrell A, Nurse P. Nuclear membrane protein Lem2 regulates nuclear size through membrane flow. Nat Commun 10, 1871 (2019).

20. Brachner A, Reipert S, Foisner R, Gotzmann J. LEM2 is a novel MAN1-related inner nuclear membrane protein associated with A-type lamins. J Cell Sci 118, 5797–5810 (2005).

21. Hirano Y, et al. Lem2 is retained at the nuclear envelope through its interaction with Bqt4 in fission yeast. Genes Cells 23, 122–135 (2018).

22. Huber MD, Guan T, Gerace L. Overlapping functions of nuclear envelope proteins NET25 (Lem2) and emerin in regulation of extracellular signal-regulated kinase signaling in myoblast differentiation. Mol Cell Biol 29, 5718–5728 (2009).

23. Cantwell H, Nurse P. A systematic genetic screen identifies essential factors involved in nuclear size control. PLoS genetics 15, e1007929 (2019).

24. Cantwell H, Nurse P. Unravelling nuclear size control. Curr Genet, (2019).

25. Jevtic P, Schibler AC, Wesley CC, Pegoraro G, Misteli T, Levy DL. The nucleoporin ELYS regulates nuclear size by controlling NPC number and nuclear import capacity. EMBO reports 20, (2019).

26. Lee JS, et al. Nuclear lamin A/C deficiency induces defects in cell mechanics, polarization, and migration. Biophys J 93, 2542–2552 (2007).

27. Wirtz D, Konstantopoulos K, Searson PC. The physics of cancer: the role of physical interactions and mechanical forces in metastasis. Nat Rev Cancer 11, 512–522 (2011).

28. Kraning-Rush CM, Califano JP, Reinhart-King CA. Cellular traction stresses increase with increasing metastatic potential. PLoS One 7, e32572 (2012).

29. Bussolati G, Maletta F, Asioli S, Annaratone L, Sapino A, Marchio C. “To be or not to be in a good shape”: diagnostic and clinical value of nuclear shape irregularities in thyroid and breast cancer. Advances in experimental medicine and biology 773, 101–121 (2014).

30. Veltri RW, Christudass CS. Nuclear morphometry, epigenetic changes, and clinical relevance in prostate cancer. Advances in experimental medicine and biology 773, 77–99 (2014).

31. Pufall MA. Glucocorticoids and Cancer. Advances in experimental medicine and biology 872, 315–333 (2015).

32. Harshbarger W, et al. Structural and Biochemical Analyses Reveal the Mechanism of Glutathione S-Transferase Pi 1 Inhibition by the Anti-cancer Compound Piperlongumine. J Biol Chem 292, 112–120 (2017).

33. Lautscham LA, et al. Migration in Confined 3D Environments Is Determined by a Combination of Adhesiveness, Nuclear Volume, Contractility, and Cell Stiffness. Biophys J 109, 900–913 (2015).

34. Davidson PM, Denais C, Bakshi MC, Lammerding J. Nuclear deformability constitutes a rate-limiting step during cell migration in 3-D environments. Cell Mol Bioeng 7, 293–306 (2014).

35. Brunner CA, Ehrlicher A, Kohlstrunk B, Knebel D, Kas JA, Goegler M. Cell migration through small gaps. Eur Biophys J 35, 713–719 (2006).

36. Ananthakrishnan R, Ehrlicher A. The forces behind cell movement. Int J Biol Sci 3, 303–317 (2007).

37. Kim JK, Louhghalam A, Lee G, Schafer BW, Wirtz D, Kim DH. Nuclear lamin A/C harnesses the perinuclear apical actin cables to protect nuclear morphology. Nat Commun 8, 2123 (2017).

38. Saleem S, Khan R, Afzal M, Kazmi I. Oxyphenbutazone promotes cytotoxicity in rats and Hep3B cellsvia suppression of PGE2 and deactivation of Wnt/beta-catenin signaling pathway. Molecular and cellular biochemistry 444, 187–196 (2018).

39. Fagone P, et al. Identification of novel chemotherapeutic strategies for metastatic uveal melanoma. Scientific reports 7, 44564 (2017).

40. Schneider NFZ, et al. Cytotoxic and cytostatic effects of digitoxigenin monodigitoxoside (DGX) in human lung cancer cells and its link to Na,K-ATPase. Biomed Pharmacother 97, 684–696 (2018).

41. Silva IT, et al. Cytotoxicity of AMANTADIG - a semisynthetic digitoxigenin derivative - alone and in combination with docetaxel in human hormone-refractory prostate cancer cells and its effect on Na(+)/K(+)-ATPase inhibition. Biomed Pharmacother 107, 464–474 (2018).

42. Bharadwaj U, et al. Drug-repositioning screening identified piperlongumine as a direct STAT3 inhibitor with potent activity against breast cancer. Oncogene 34, 1341–1353 (2015).

43. Raj L, et al. Selective killing of cancer cells by a small molecule targeting the stress response to ROS. Nature 475, 231–234 (2011).

44. Jeong CH, et al. Piperlongumine Induces Cell Cycle Arrest via Reactive Oxygen Species Accumulation and IKKbeta Suppression in Human Breast Cancer Cells. Antioxidants (Basel) 8, (2019).

45. Zhang Q, et al. Piperlongumine, a Novel TrxR1 Inhibitor, Induces Apoptosis in Hepatocellular Carcinoma Cells by ROS-Mediated ER Stress. Front Pharmacol 10, 1180 (2019).

46. Zhou L, Li M, Yu X, Gao F, Li W. Repression of Hexokinases II-Mediated Glycolysis Contributes to Piperlongumine-Induced Tumor Suppression in Non-Small Cell Lung Cancer Cells. Int J Biol Sci 15, 826–837 (2019).

47. Liu Z, et al. Piperlongumine-induced nuclear translocation of the FOXO3A transcription factor triggers BIM-mediated apoptosis in cancer cells. Biochem Pharmacol 163, 101–110 (2019).

48. Mgbeahuruike EE, Stalnacke M, Vuorela H, Holm Y. Antimicrobial and Synergistic Effects of Commercial Piperine and Piperlongumine in Combination with Conventional Antimicrobials. Antibiotics (Basel) 8, (2019).

49. Wang Y, et al. Discovery of piperlongumine as a potential novel lead for the development of senolytic agents. Aging (Albany NY) 8, 2915–2926 (2016).

50. Sanchez C, Reines EH, Montgomery SA. A comparative review of escitalopram, paroxetine, and sertraline: Are they all alike? Int Clin Psychopharmacol 29, 185–196 (2014).

51. Then CK, Liu KH, Liao MH, Chung KH, Wang JY, Shen SC. Antidepressants, sertraline and paroxetine, increase calcium influx and induce mitochondrial damage-mediated apoptosis of astrocytes. Oncotarget 8, 115490–115502 (2017).

52. Havercroft JC, Quinlan RA, Gull K. Binding of parbendazole to tubulin and its influence on microtubules in tissue-culture cells as revealed by immunofluorescence microscopy. J Cell Sci 49, 195–204 (1981).

53. Quinlan RA, Roobol A, Pogson CI, Gull K. A correlation between in vivo and in vitro effects of the microtubule inhibitors colchicine, parbendazole and nocodazole on myxamoebae of Physarum polycephalum. J Gen Microbiol 122, 1–6 (1981).

54. Kainuma K, et al. beta2 adrenergic agonist suppresses eosinophil-induced epithelial-to-mesenchymal transition of bronchial epithelial cells. Respir Res 18, 79 (2017).

55. Zhang P, et al. Post-radiotherapy maintenance treatment with fluticasone propionate and salmeterol for lung cancer patients with grade III radiation pneumonitis: A case report. Medicine (Baltimore) 97, e10681 (2018).

56. Tuglu MM, Bostanabad SY, Ozyon G, Dalkilic B, Gurdal H. The role of dualspecificity phosphatase 1 and protein phosphatase 1 in beta2adrenergic receptormediated inhibition of extracellular signal regulated kinase 1/2 in triple negative breast cancer cell lines. Mol Med Rep 17, 2033–2043 (2018).

57. Wnorowski A, et al. Concurrent activation of beta2-adrenergic receptor and blockage of GPR55 disrupts pro-oncogenic signaling in glioma cells. Cell Signal 36, 176–188 (2017).

58. Rivero EM, et al. The beta 2-Adrenergic Agonist Salbutamol Inhibits Migration, Invasion and Metastasis of the Human Breast Cancer MDA-MB-231 Cell Line. Curr Cancer Drug Targets 17, 756–766 (2017).

59. Lu Y, et al. Isoprenaline/beta2-AR activates Plexin-A1/VEGFR2 signals via VEGF secretion in gastric cancer cells to promote tumor angiogenesis. BMC Cancer 17, 875 (2017).

60. Skaga E, Skaga IO, Grieg Z, Sandberg CJ, Langmoen IA, Vik-Mo EO. The efficacy of a coordinated pharmacological blockade in glioblastoma stem cells with nine repurposed drugs using the CUSP9 strategy. J Cancer Res Clin Oncol 145, 1495–1507 (2019).

61. Tian YS, Chen KC, Zulkefli ND, Maner RS, Hsieh CL. Evaluation of the Inhibitory Effects of Genipin on the Fluoxetine-Induced Invasive and Metastatic Model in Human HepG2 Cells. Molecules 23, (2018).

62. Chen WT, Hsu FT, Liu YC, Chen CH, Hsu LC, Lin SS. Fluoxetine Induces Apoptosis through Extrinsic/Intrinsic Pathways and Inhibits ERK/NF-kappaB-Modulated Anti-Apoptotic and Invasive Potential in Hepatocellular Carcinoma Cells In Vitro. Int J Mol Sci 20, (2019).

63. Davidson PM, Sliz J, Isermann P, Denais C, Lammerding J. Design of a microfluidic device to quantify dynamic intra-nuclear deformation during cell migration through confining environments. Integr Biol (Camb) 7, 1534–1546 (2015).

64. Keys J, Windsor A, Lammerding J. Assembly and Use of a Microfluidic Device to Study Cell Migration in Confined Environments. Methods Mol Biol 1840, 101–118 (2018).

65. Elacqua JJ, McGregor AL, Lammerding J. Automated analysis of cell migration and nuclear envelope rupture in confined environments. PLoS One 13, e0195664 (2018).

