## Supplemental Figures and Supplemental Table legends for "Chemical-Genetic Interrogation of Nuclear Size Control Reveals Cancer-Specific Effects on Cell Migration and Invasion"

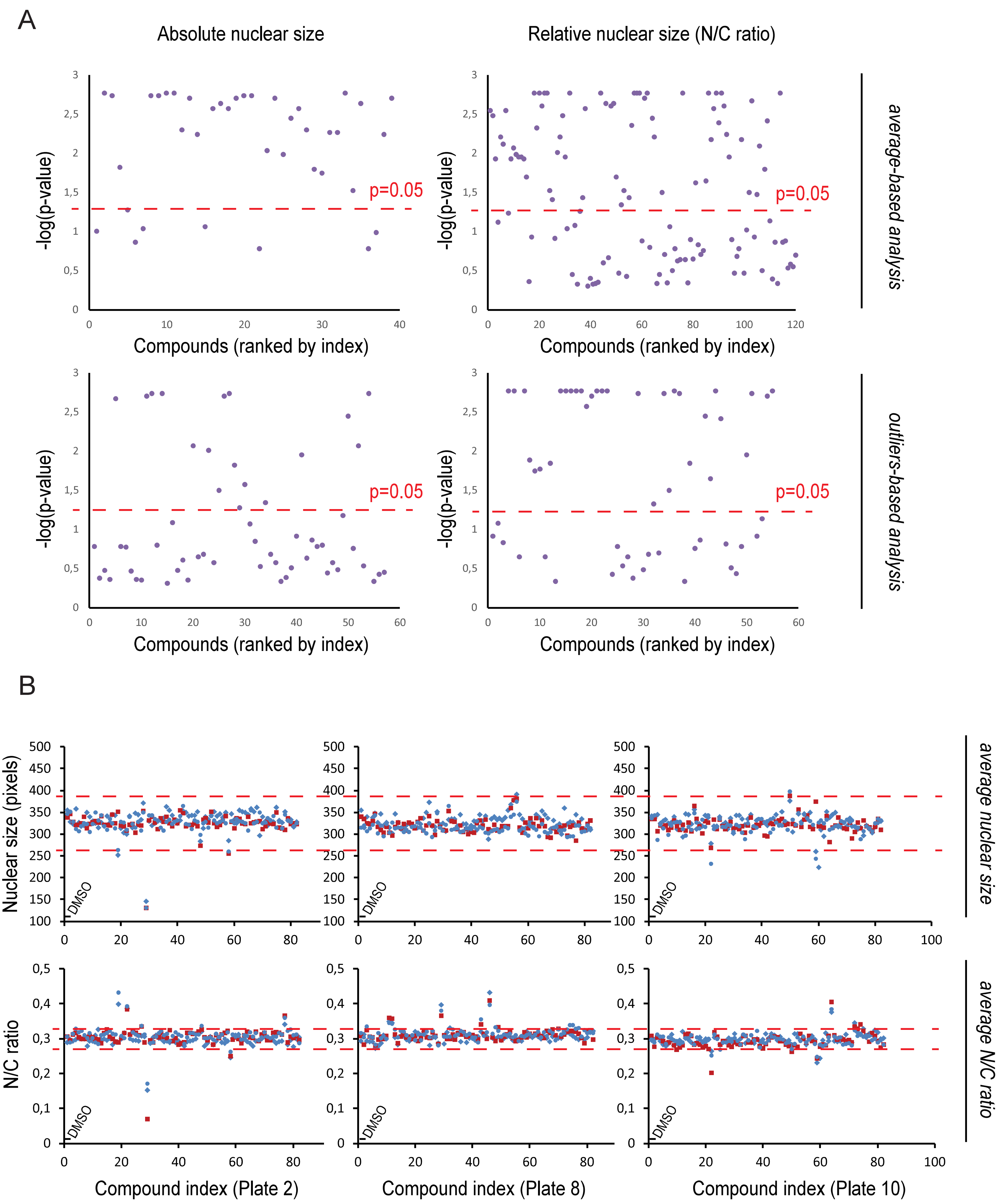
A
Absolute nuclear size
Relative nuclear size (N/C ratio)
-log(p-value)
p=0.05
Compounds (ranked by index)
average-based analysis
outliers-based analysis
B
Nuclear size (pixels)
N/C ratio
DMSO
average nuclear size
average N/C ratio
Compound index (Plate 2)
Compound index (Plate 8)
Compound index (Plate 10)

**Supplementary Figure S1. Screen data reproducibility.**

A. Scatter plots showing the Wilcoxon rank tests p-values (log scale, vertical axis) for each "hit" compound (horizontal axis) compared with DMSO across all replicates (see Methods). Tests were performed from the average nuclear size (left scatters) and N/C ratio metrics (right scatters) data acquired on PC3 cells with 6h treatment (3 replicates), and using average-based (top scatters) and outliers-based (bottom scatters) analyses. The significance level (p=0.05) is shown as red dashed lines. B. Examples of raw data scatter plots showing the average nuclear size (top scatters) and average N/C ratio (bottom scatters) as directly output by our Matlab analysis scripts for compound plate 2 (left), plate 8 (middle) and plate 10 (right). Blue circle, blue diamond and red squares represent the 3 replicates (PC3 cells, 6h treatment). For each plate/replicate/metric, the first 2 dots of the scatter represent the value metric averaged over all in-plate DMSO control wells (16 wells), and compounds are then listed from A2, A3 … A11, B2 … to H11. Dashed lines indicate the hit detection thresholds for each metric, as determined for PC3 cells, 6h treatment (see Methods).

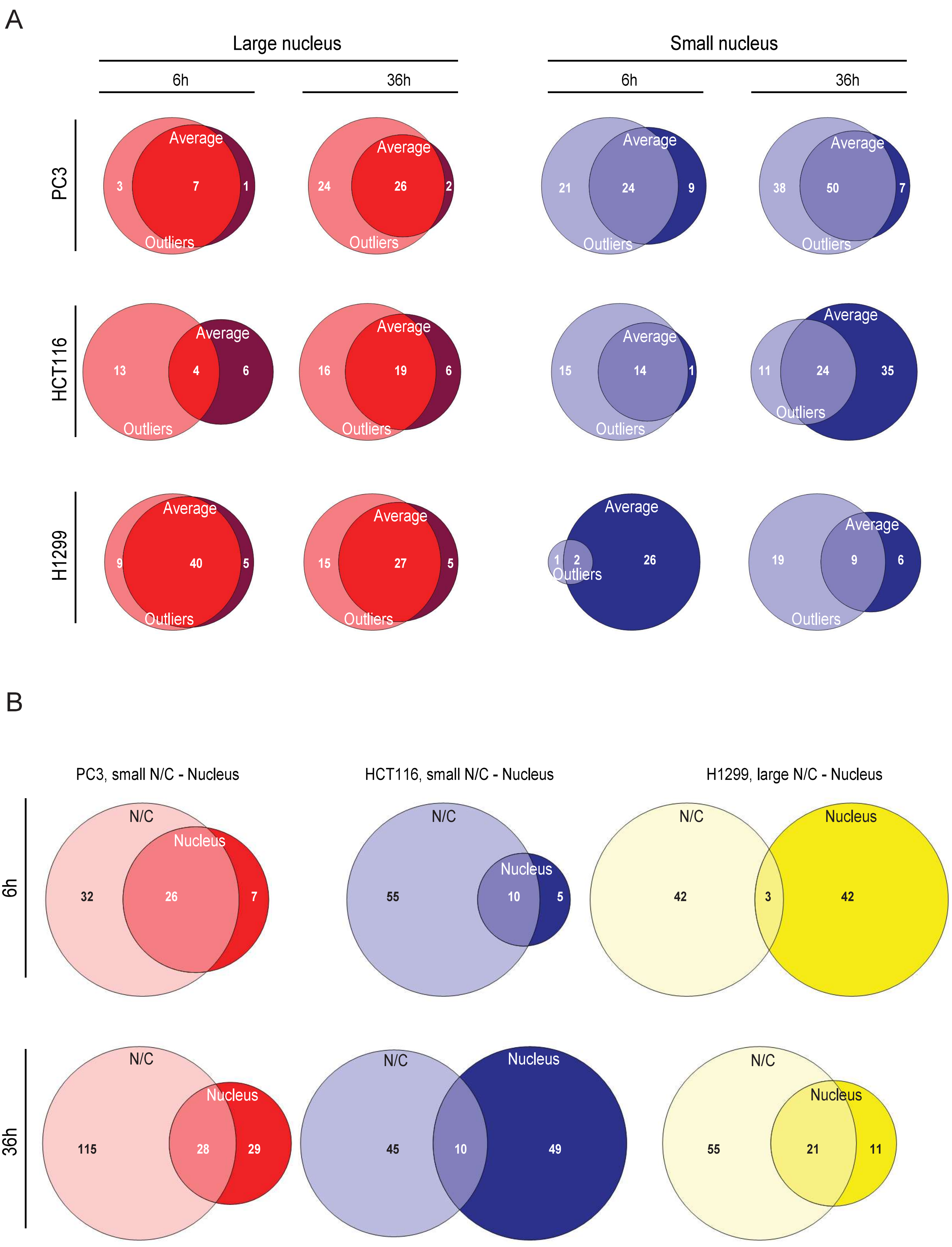

A
Large nucleus
Small nucleus
6h
36h
6h
36h
PC3
Average
Outliers
3
7
1
24
26
2
21
24
9
38
50
7
HCT116
13
4
6
16
19
6
15
14
1
11
24
35
H1299
9
40
5
15
27
5
1
2
26
19
9
6
B
PC3, small N/C - Nucleus
HCT116, small N/C - Nucleus
H1299, large N/C - Nucleus
6h
36h
N/C
Nucleus
32
26
7
55
10
5
42
3
42
115
28
29
45
10
49
55
21
11

**Supplementary Figure S2. Analysis methodology, but not the choice of metric, yielded overlapping results.**

A. Venn diagrams showing the overlap in compounds increasing (left diagrams, red) or reducing (right diagrams, blue) the absolute nuclear size, as determined by average-based (dark colours) or outlier-based (light colours) strategies as indicated. Overlaps in analysis methodologies were computed for PC3 (top), HCT116 (middle) and H1299 (bottom) cells treated for 6h or 36 h as indicated. B. Venn diagrams showing the overlap in compounds correcting cancer-related nuclear changes as determined using the average nuclear size (dark colours) vs average N/C ratio (light colours). Overlaps in hits determined using the two metrics were computed separately for PC3 (red), HCT116 (blue) and H1299 (yellow) cells treated for 6h (top row) or 36 h (bottom row) as indicated.

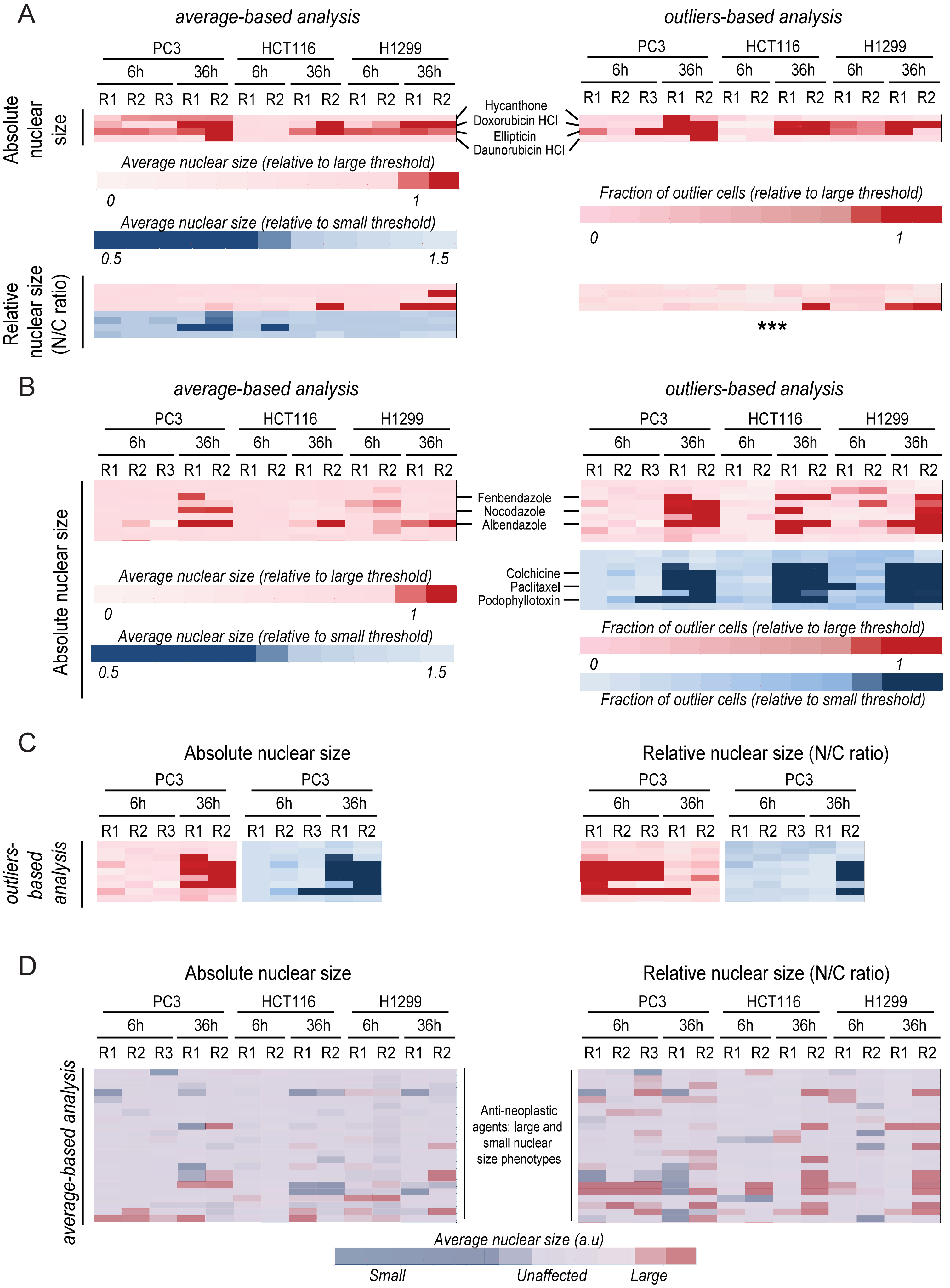
A
average-based analysis
outliers-based analysis
PC3
HCT116
H1299
6h
36h
R1 R2 R3 R1 R2 R1 R2 R1 R2 R1 R2 R1 R2
Absolute nuclear size
Hycanthone
Doxorubicin HCl
Ellipticin
Daunorubicin HCl
Average nuclear size (relative to large threshold)
0
1
Average nuclear size (relative to small threshold)
0.5
1.5
Fraction of outlier cells (relative to large threshold)
Relative nuclear size (N/C ratio)
***
B
Absolute nuclear size
Fenbendazole
Nocodazole
Albendazole
Colchicine
Paclitaxel
Podophyllotoxin
Fraction of outlier cells (relative to small threshold)
C
Relative nuclear size (N/C ratio)
outliers-based analysis
D
average-based analysis
Anti-neoplastic agents: large and small nuclear size phenotypes
Average nuclear size (a.u)
Small
Unaffected
Large

**Supplementary Figure S3. Comparison of absolute and relative nuclear size responses to different treatment duration informs on compound mechanism of action.** A. Heat maps showing nuclear size data (top heat maps) and the N/C ratio data (bottom heat maps) across tumour type/replicates (horizontal) treated with DNA intercaling agents (vertical), as determined by average-based analysis (left) and outliers-based analysis (right). Both averages, and the fractions of outlier cells for nuclear size and N/C ratio metrics were normalized to the detection thresholds for large/small nucleus or N/C ratio (see Methods), and colour-coded as indicated. Darker colours correspond to stronger phenotypes. Data for the fraction of outlier cells with abnormally small N/C ratio couldn't be determined due to renormalization issues (***, see Methods). B. Heat maps showing absolute nuclear size across tumour type/replicates (horizontal) treated with microtubule polymerization inhibitors (vertical), as determined by outliers-based analysis (right) and average-based analysis (left). Data normalization and colour-coding were performed similarly to panel (A). The nine microtubule polymerization inhibitor compounds are the same on the three heat maps and listed in the same order, particular compounds names are indicated next to different heat maps for convenience. C. Heat maps showing absolute nuclear size (left) and relative nuclear size (N/C ratio, right) across PC3 replicates (horizontal) treated with microtubule polymerization inhibitors (vertical), as determined by outliers-based analysis. Fractions of outlier cells data was normalized and colour-coded as indicated on panel B. D. Heat maps showing the mean nuclear size (left) and the mean N/C ratio (right) across tumour type/replicates (horizontal) treated with all Prestick compounds documented as antineoplastic. Nuclear size and N/C ratio have been normalized to both the detection thresholds for large (yielding a red heat map) and for small nucleus (yielding a blue heat map), both of which have been colour-coded as indicated. Then, both red and blue heat maps have been superimposed to yield the multi-colour heat map shown. Almost all antineoplastic agents affect the absolute nuclear size and/or the N/C ratio in at least one cell line/condition.

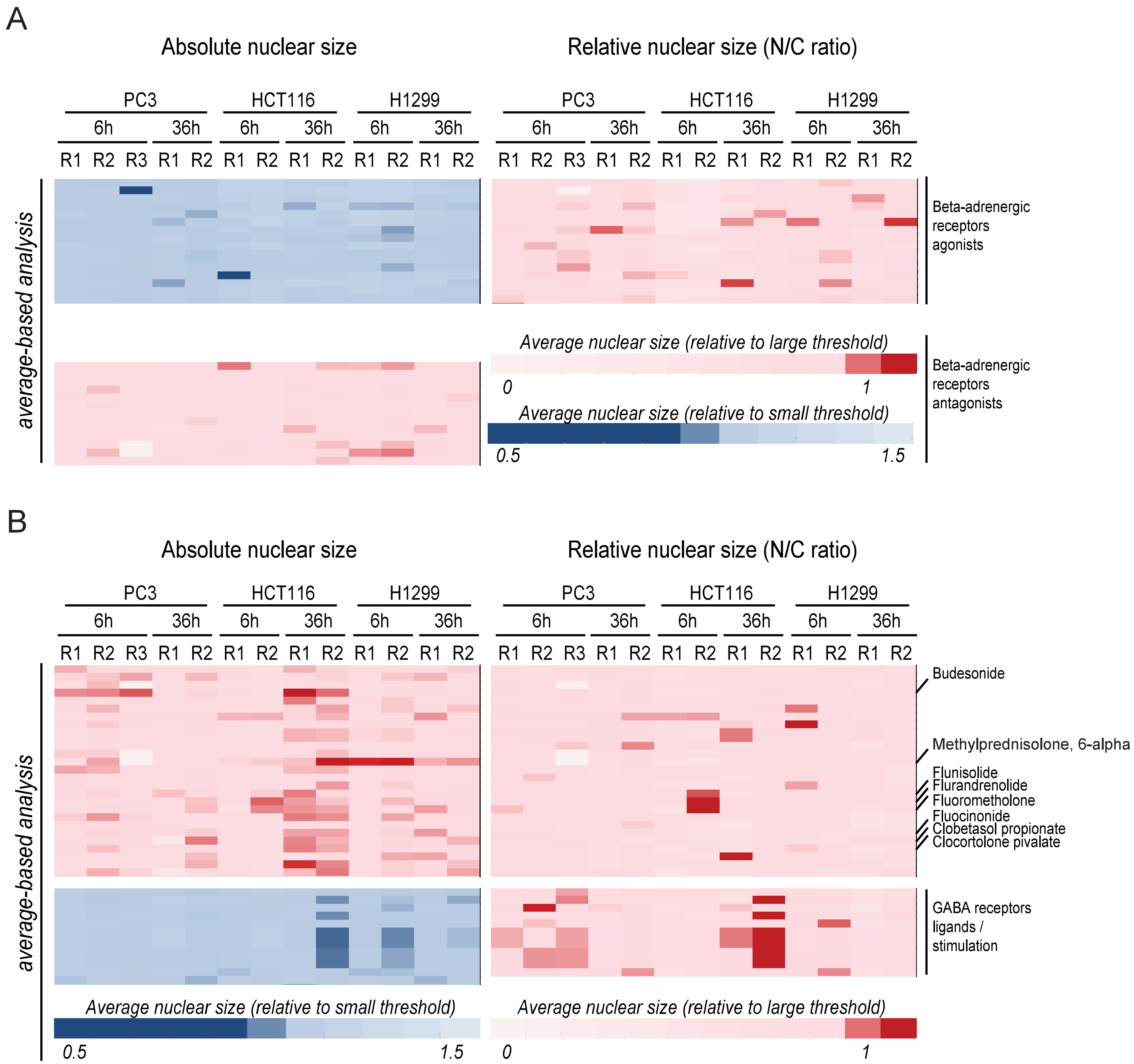
A
Absolute nuclear size
Relative nuclear size (N/C ratio)
PC3
HCT116
H1299
6h
36h
R1 R2 R3 R1 R2 R1 R2 R1 R2 R1 R2 R1 R2
average-based analysis
Beta-adrenergic receptors agonists
Beta-adrenergic receptors antagonists
Average nuclear size (relative to large threshold)
0
1
Average nuclear size (relative to small threshold)
0.5
1.5
B
Absolute nuclear size
Relative nuclear size (N/C ratio)
Budesonide
Methylprednisolone, 6-alpha
Flunisolide
Flurandrenolide
Fluorometholone
Fluocinonide
Clobetasol propionate
Clocortolone pivalate
GABA receptors ligands / stimulation
Average nuclear size (relative to small threshold)
Average nuclear size (relative to large threshold)

**Supplementary Figure S4. On the importance of knowing which of absolute or relative nuclear size actively contributes to metastasis.**

A. Heat maps showing the mean nuclear size (left) and the mean N/C ratio (right) across tumour type/replicates (horizontal) treated with beta-adrenergic receptor agonists (top heat maps) or beta-adrenergic receptor antagonists (bottom heat map). Both heat maps show particular regions from Figures 3-4, magnified for better visualization. Nuclear size and N/C ratio were normalized to the detection thresholds for large nucleus (red heat maps, see Methods), or for small nucleus (blue heat maps), and colour-coded as indicated. B. Heat maps showing the mean nuclear size (left) and the mean N/C ratio (right) across tumour type/replicates (horizontal) treated with glucocorticoids (top heat maps) or GABA receptor ligands/stimulators (bottom heat map). Nuclear size and N/C ratio were normalized to the detection thresholds for large nucleus or N/C (red heat maps, see Methods), or for small nucleus or N/C (blue heat maps), and colour-coded as indicated. Some compounds of particular interest are indicated on the side.

A

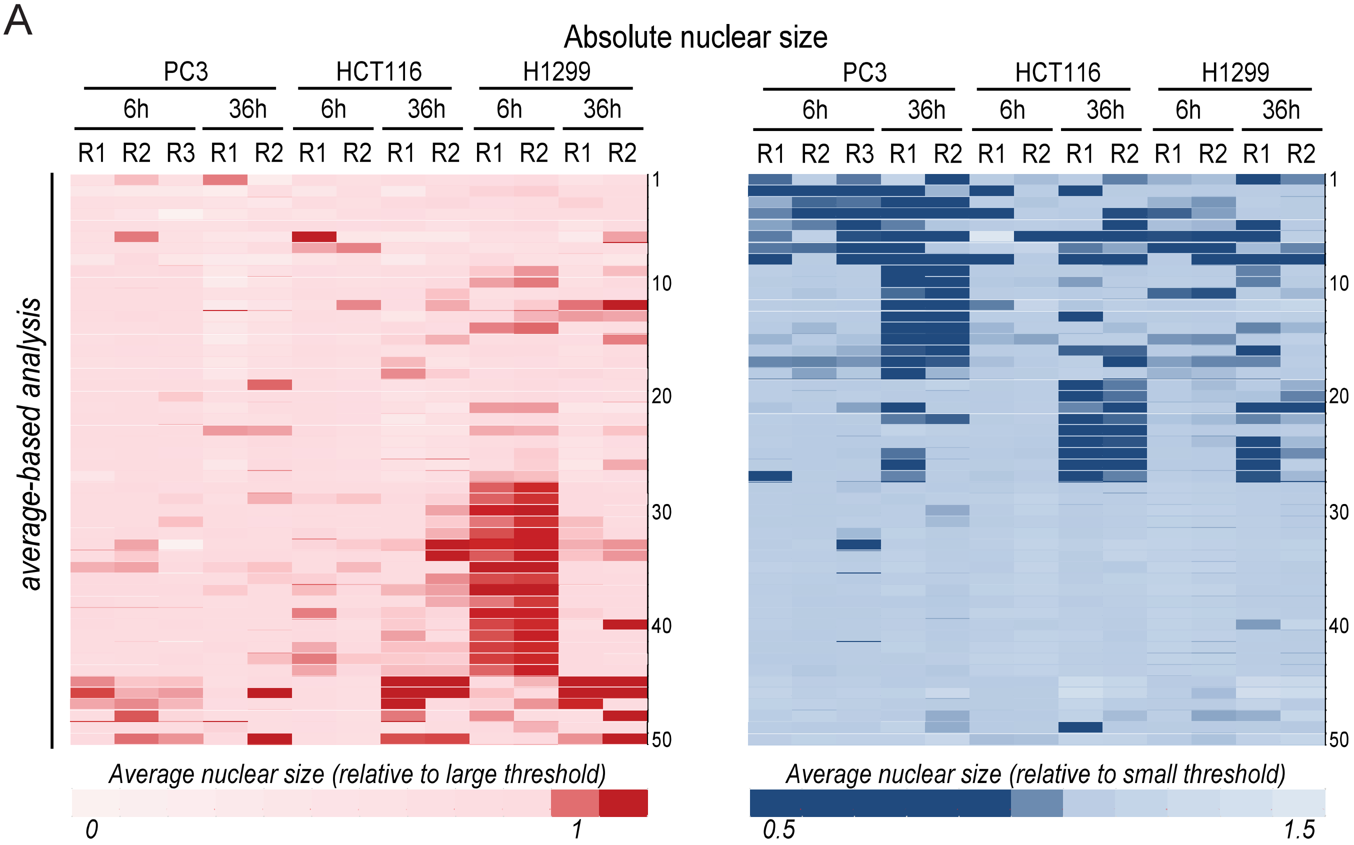


| | | | | | | | |
|---|---|---|---|---|---|---|---|
| 1 | Disulfiram | 14 | Emetine di-HCl | 27 | Azacytidine 5 | 40 | Proscillaridin A |
| 2 | Tomatine | 15 | Piperlongumine | 28 | Hydroflumethiazide | 41 | Etilefrin HCl |
| 3 | Astemizole | 16 | Lycorine HCl | 29 | Metaraminol bitartrate | 42 | Letrozole |
| 4 | Perhexiline maleate | 17 | Sertraline | 30 | Flufenamic acid | 43 | (+,-)-Synephrine |
| 5 | Ebselen | 18 | Paroxetin HCl | 31 | Trimethoprim | 44 | Verteporfin |
| 6 | Alexidine di-HCl | 19 | Danazol | 32 | Fenspiride HCl | 45 | Monobenzone |
| 7 | Cantharidin | 20 | Griseofulvin | 33 | Methylprednisolone 6-α | 46 | Trifluridine |
| 8 | Oxyphenbutazone | 21 | Puromycin di-HCl | 34 | Debrisoquin sulfate | 47 | Antimycin A |
| 9 | Anisomycin | 22 | Resveratrol | 35 | Harmol HCl mono-H2O | 48 | Mitoxantrone di-HCl |
| 10 | Roxatidine acetate | 23 | Cilostazol | 36 | Metoxy-6-harmalan | 49 | Parbendazole |
| 11 | Metixene HCl | 24 | Eburnamonine | 37 | Levonordefrin | 50 | Camptothecine (S,+) |
| 12 | Parthenolide | 25 | Monensin Na salt | 38 | N-acetyl-L-leucine | | |
| 13 | Prenylamine lactate | 26 | Dienestrol | 39 | (+)-Isoproterenol bitartrate | | |

B

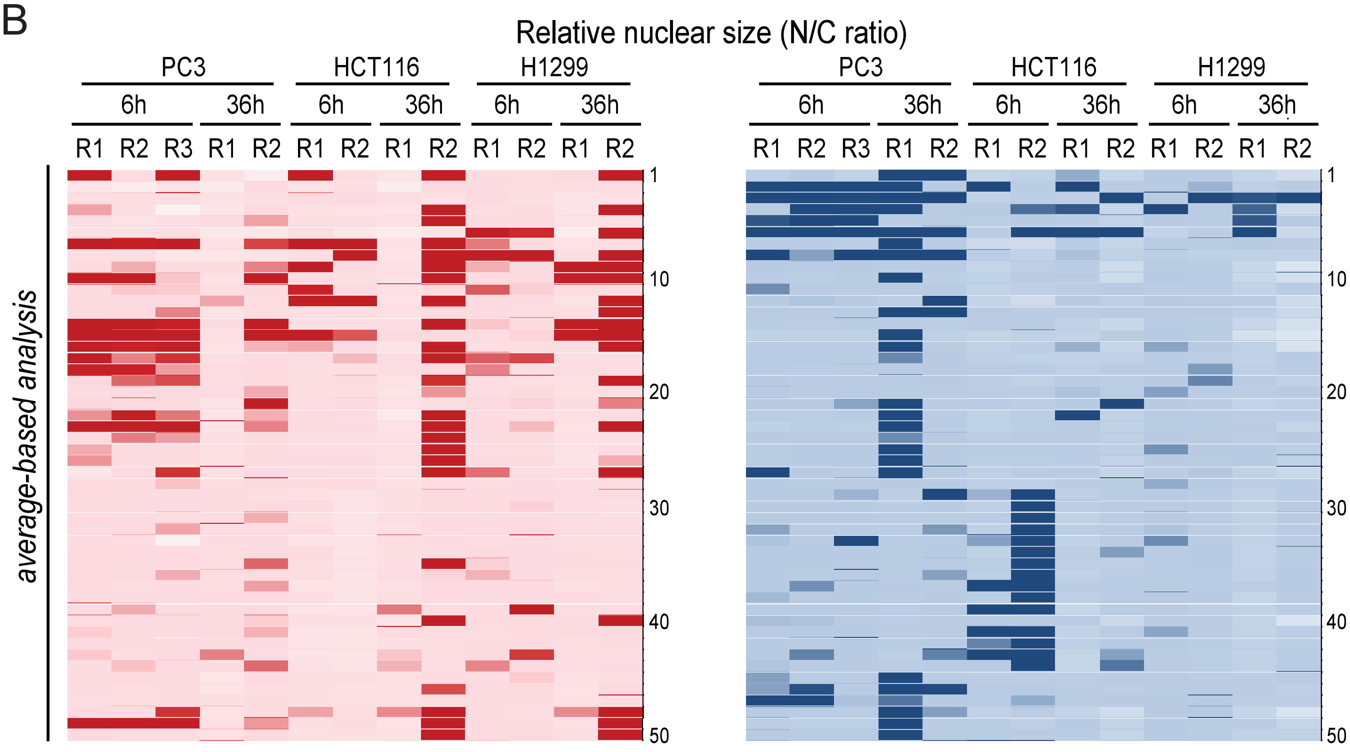

**Supplementary Figure S5. Individual compounds spanning many pharmacological classes can correct cancer-associated nuclear size changes.**

A. Heat maps showing the mean nuclear size (A) and the mean N/C ratio (B) across tumour type/replicates (horizontal) treated with the Prestwick compounds (individual rows) that showed the strongest effects in one or several conditions (as listed in Table 1). For clarity, compounds are labeled with numbers on heat maps and listed in the Table shown on panel (A). Cell population-averaged nuclear size and N/C ratio have been normalized to both the detection thresholds for large (yielding a red heat map) and for small nucleus (yielding a blue heat map), and colour-coded as indicated.

**Supplementary Table 1.**

Summary table showing the fraction of hits replicated once, twice or three times, using the two metrics and two analysis strategies introduced in the main text, and that yielded a p-value lower than 0.05 upon a compound vs DMSO Wilcoxon rank-test.

**Supplementary Table 2.**

Effects across cell lines and duration of treatment of compounds identified as affecting the average absolute nuclear size in at least one condition.

**Supplementary Table 3.**

Effects across cell lines and duration of treatment of compounds identified as affecting the average N/C ratio in at least one condition.

**Supplementary Table 4.**

Effects across cell lines and duration of treatment of compounds identified as affecting the fractions of outlier cells with abnormally large or small nucleus in at least one condition.

**Supplementary Table 5.**

Effects across cell lines and duration of treatment of compounds identified as affecting the fractions of outlier cells with abnormally large or small N/C ratio in at least one condition.
