## Supplemental Table 1 for "Chemical-Genetic Interrogation of Nuclear Size Control Reveals Cancer-Specific Effects on Cell Migration and Invasion"

| **Metric** | **Analysis strategy** | **% of 1-rep hits with p<0.05** | **% of 2-rep hits with p<0.05** | **% of 3-rep hits with p<0.05** |
| --- | --- | --- | --- | --- |
| Nuclear size | Average | 80,6 | 66,7 | 100 |
| Nuclear size | Outliers | 16,3 | 87,5 | 100 |
| N/C ratio | Average | 40 | 93,3 | 100 |
| N/C ratio | Outliers | 38,6 | 88,9 | 100 |
